# Membrane composition influences the conformation and function of the dopamine transporter *in vivo*

**DOI:** 10.1101/755819

**Authors:** Wendy M. Fong, Kevin Erreger, Se Joon Choi, India Reddy, Christopher W. Johnson, Eugene V. Mosharov, Jonathan A. Javitch, Ai Yamamoto

## Abstract

The biophysical and biochemical properties of membrane lipids can alter the conformation and function of membrane-spanning proteins, yet the specific, physiological consequence *in vivo* of changing the membrane milieu for a specific protein has been rarely investigated. Using various genetic approaches to eliminate expression of the membrane-associated protein Flotillin-1, we have found that the lipid environment of the dopamine transporter (DAT) is necessary for mice to respond to amphetamine but not cocaine, because the localization of DAT to cholesterol-rich membranes is required for a DAT conformation that is essential for reverse transport of dopamine. Furthermore, a conditional rather than constitutive loss-of-function approach was necessary to reveal this phenotype, indicating a broader role for membrane-protein interactions that are modulated by Flotillin-1. Taken together, these findings demonstrate how interaction of a transmembrane protein with its membrane environment can regulate distinct events in the vertebrate brain that give rise to specific behavioral outcomes.

## Introduction

The biophysical and biochemical properties of membrane lipids modulate the conformation and function of membrane-spanning proteins, and thereby influence diverse cellular functions. Whether this occurs through the formation of discrete protein-lipid microdomains or by other mechanical means such as membrane deformation, is an open question {Simons, 2011 #5262; Sezgin, 2017 #5263}. Fundamentally however, it has been difficult to test the specific, physiological consequences following a change in the membrane milieu for a given protein.

Membrane-associated proteins that are members of the SPFH family, such as Flotillin-1/Reggie-2 (Flot1) is linked to a wide range of essential proteins from cell adhesion molecules to receptors (Banning et al., 2011; Bodin et al., 2014; Cremona et al., 2011; Otto and Nichols, 2011; Stuermer, 2011, 2012; Stuermer and Plattner, 2005), and is hypothesized to scaffold these substrates into cholesterol-rich membrane domains to facilitate their function. Members of the SLC6 family of cell surface transporters, such as the dopamine transporter (DAT), have been shown to partially partition to detergent-resistant membranes (DRMs) upon homogenization with 1% Brij58 (Adkins et al., 2007; Cremona et al., 2011; Hong and Amara, 2010; Jones et al., 2012; Magnani et al., 2004).

DAT is responsible for uptake and clearance of extracellular dopamine (DA) from the perisynaptic space, and thus modulation of DAT activity can influence a wide range of DA-mediated behaviors from movement to cognition (Jaber et al., 1997). Although the partitioning of DAT and other transporters into cholesterol-rich membranes represented by DRMs has been replicated by several groups (Cremona et al., 2011; Hong and Amara, 2010; Jones et al., 2012), the larger significance of this partitioning remains unclear. By super-resolution microscopy, DAT has been found associated to cholesterol-enriched, Flot1 positive nanodomains (Rahbek-Clemmensen et al., 2017), that is well correlated with the biochemical data. Flot1 depletion in cell lines has no impact on DAT-mediated uptake of DA, but in primary mesencephalic neurons, Flot1 depletion diminishes the non-exocytic release of DA through DAT after administration of the psychostimulant amphetamine (AMPH) (Cremona et al., 2011), and diminished AMPH-evoked locomotion in *Drosophila* larvae heterologously expressing human DAT (hDAT)(Pizzo et al., 2013). These data suggest that Flot1 is required for the reverse transport of DA through DAT, also known as DAT efflux, and that efflux contributes towards the increased perisynaptic levels of DA *in vivo* in response to AMPH (Sitte and Freissmuth, 2015). The extent to which reverse transport contributes to the behavioral outcomes of acute AMPH in vertebrates has also been questioned (Daberkow et al., 2013; Siciliano et al., 2014).

To test the physiological importance of the relationship between Flot1 and DAT, we created a series of genetically modified mice in which we conditionally or constitutively removed Flot1. We find that DAT partitions into cholesterol-rich DRMs in a Flot1-dependent manner, and DAT partitioning increases or decreases upon the administration of amphetamine (AMPH) or cocaine (COC), respectively. Loss of Flot1 in DAergic neurons prevents DAT from partitioning to buoyant fractions, but leads to no change in basal DA neuron parameters, including DA uptake by DAT, indicating that the partitioning does not reflect the direct interaction of DAT with cholesterol that is necessary for DAT function. Moreover, Flot1 was required for mice to mount a normal response to AMPH, but not COC, clearly implicating distinct modes of action for these two well-studies psychostimulants. Mechanistic studies revealed that DAT partitioning stabilizes a conformation of DAT that interacts with the membrane phospholipid PIP_2_, which is essential for the reverse transport of DA through DAT. Notably, the blunted response to AMPH- and Flot1-dependent partitioning of DAT was observed under conditional but not constitutive elimination of Flot1, indicating that the latter might evoke epigenetic events that mask the importance of Flot1-mediated scaffolding. These data indicate that the membrane environment of DAT, which is governed by Flot1, is important for AMPH-induced reverse transport of DA because a cholesterol-rich environment promotes a conformation of DAT that is favorable for DA efflux.

## Results

### Flot1 is required in DA neurons for DAT partitioning into cholesterol-rich membranes upon administration of AMPH

To determine how Flot1 expression might influence the ability of mice to respond to the psychostimulant AMPH (Cremona et al., 2011), we created a conditional KO mouse that genetically eliminates Flot1 in DAergic neurons. Mice carrying a conditional *Flot1* allele (Flot1^fl/fl^) were crossed to mice carrying the DAT^iresCre^ allele, resulting in a Flot1 conditional knockout (Flot1 cKO) strain (Figures S1A-C). Unlike DAT KO mice (Giros et al., 1996), but similar to previously published Flot1 KO mice (Banning et al., 2011; Bitsikas et al., 2014; Bodrikov et al., 2017; Ludwig et al., 2010), Flot1 cKO mice demonstrate no overt phenotype and were able to breed and thrive comparably to littermate controls. Given the report of modestly reduced DAT levels in the heterozygous DAT^iresCre^ mice (Backman et al., 2006), mice heterozygous for conditional deletion of *Flot1* (Flot1 cHet) are used as control.

To examine DAT partitioning, we dissected striata from Flot1 cHet and cKO mice in the presence of absence of AMPH and isolated 1% Brij58-resistant DRMs. Immunoblotting revealed that, similar to heterologously expressed DAT, endogenous DAT partitions into DRMs in a Flot1-dependent manner which becomes more robust upon administration of AMPH (Figure 1A and B). Pre-treatment of the lysates with the cholesterol chelator methyl-beta-cyclodextrin (MβC) confirmed previous reports that DAT partitioning also depends on cholesterol (Figure S1D). Taken together these data suggest that the partitioning of DAT into 1% Brij58-resistant DRMs increases in response to AMPH.

**Figure 1.**
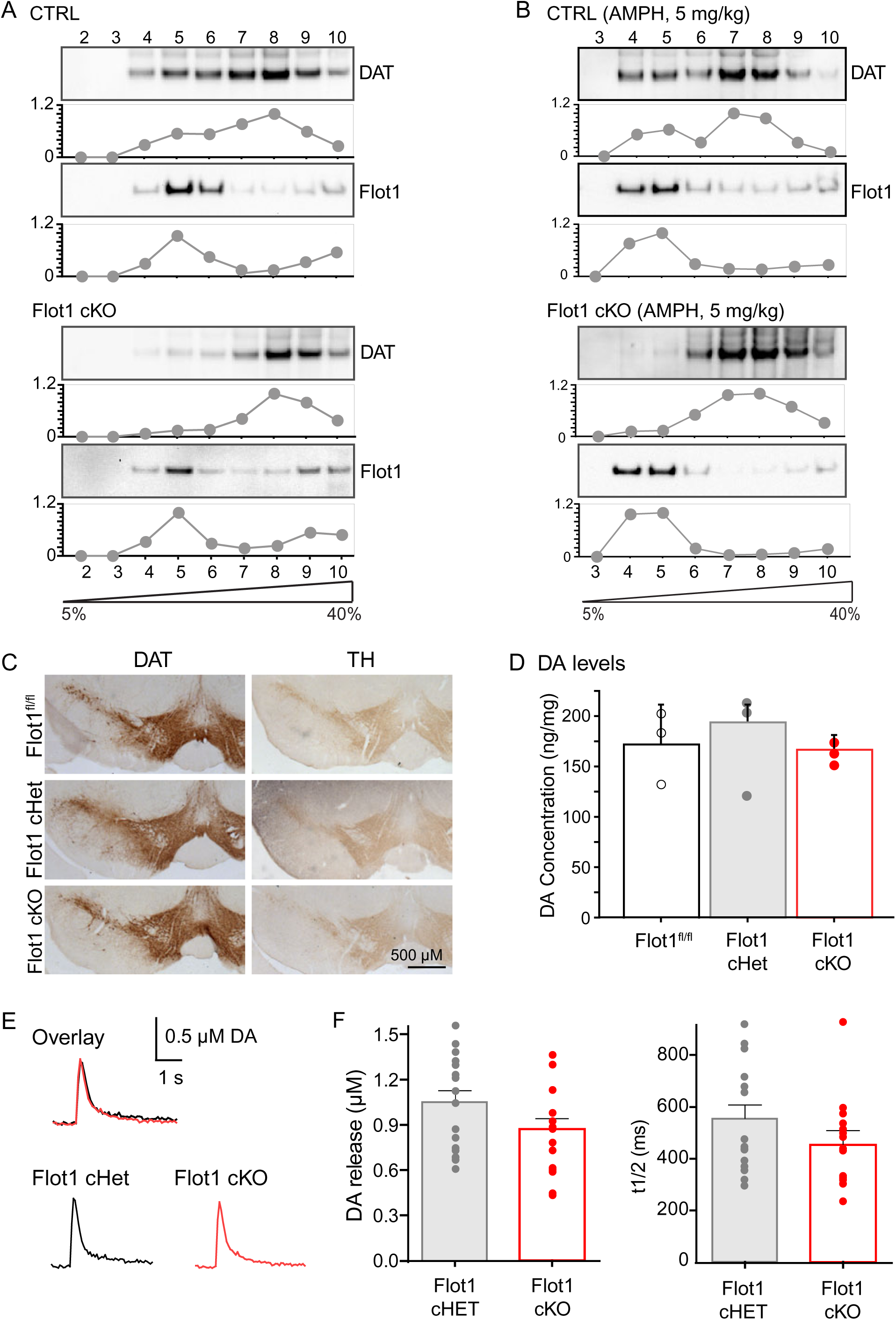
Flot1 is required to partition DAT into DRMs, but not for basal DA neuron function. **A, B.** Sucrose density gradient centrifugation reveals a differential distribution of DAT in the Flot1 cKO brains. Striatal homogenates in the absence (**A**) or presence of 5 mg/kg AMPH i.p. (**B**) were lysed in 1% Brij58 were subject to a step gradient of 5% to 40% sucrose, spun at 43,200 rpm for 20 hrs at 4°C then collected as 10 equal fractions. DAT detected in lysates from Flot1^fl/fl^ striata partitions into Flot1-positive detergent resistant membrane fractions (DRMs) that are cholesterol-dependent, as revealed by methyl-beta-cyclodextrin treatment (Figure S1D). In contrast, DAT from lysates of Flot1 cKO striata no longer partitions into DRMs. The presence of Flot1 is derived from the cast majority of the cells in the striatum that are not DAergic. **C.** Immunohistochemistry against DAT and tyrosine hydroxylase (TH) of substantia nigra (SN) from coronal brain sections from Flot1^fl/fl^, cHet, and cKO littermates. Striatum can be found in Figure S1. n = 3/genotype. **D.** HPLC reveals normal DA content in the striata of Flot1 cKO mice. DOPAC as well as norepinephrine and 5-HT are similarly unchanged (not shown). ANOVA revealed no effect of genotype on DA levels (F_(2,6)_ = 0.868, p = 0.4667)). n = 3/genotype. Data represented as mean ± S.E. with individual data points shown. **E, F**. Cyclic voltammetry in striatal slice preparations also found no difference in basal DA release and DAT-mediated uptake between cHet and cKO mice. **E.** Representative traces from cKO and cHet mice. **F.** Quantification of DA release and DA uptake as represented by t_1/2_. No significant difference was observed in either measure. (Mann Whitney U test revealed no effect of genotype on DA release (p = 0.137) or on t_1/2_ (p = 0.147)). N = 6 brain, n = 16 slices/genotype. Data represented as mean ± S.E. with individual data points shown.

### Loss of Flot1 in DA neurons does not affect basal neuron function

Structural data indicate that DAT interacts with a molecule of cholesterol to permit proper folding (Penmatsa et al., 2013), and that its association to cholesterol is essential for DAT function, especially neurotransmitter reuptake (Hong and Amara, 2010; Jones et al., 2012; Zeppelin et al., 2018)}. Given that it is unclear whether the essential association of cholesterol to DAT is associated to its Flot1-dependent partitioning, we examined whether the loss of Flot1 disrupts basal parameters of the DAergic system. Immunohistochemistry (IHC) against DAT and tyrosine hydroxylase (TH) revealed no gross neuroanatomical differences in the striatum (Figure S1E) or substantia nigra (Figure 1C), suggesting that levels of DAT and DA are unaffected by Flot1 expression levels. HPLC analyses of striatal lysates confirmed that DA levels were unchanged (Figure 1D). Given that IHC does not reflect levels of functional DAT, we also examined basal DA release and uptake dynamics in slice preparations from dorsolateral striatum by single-pulse electrical stimulation. As shown in Figures 1E and 1F, no significant difference in evoked DA release or DA uptake as measured by t_1/2_ were observed across genotype, indicating that cell surface levels of DAT are unchanged by loss of Flot1 as expected by previous studies (Block et al., 2015) and that the loss of partitioning does not interfere with the ability of DAT to interact with cholesterol. The latter findings also strongly suggest that functional DAT levels are similar across genotypes. Thus, these data indicates that the Flot1-mediated partitioning we observe is independent of this critical interaction of DAT and cholesterol, and that its loss in Flot1 cKO mice does not affect basal parameters of the DAergic system.

### Conditional loss of Flot1 in dopaminergic neurons leads to a diminished response to AMPH in mice

Given that basal DAT function does not require Flot1, we next examined the response of Flot1 cKO mice to two frequently studied psychostimulants that act on DAT: cocaine (COC) and amphetamine (AMPH). Although both compounds increase the levels of perisynaptic DA, thereby leading to behaviors such as hyper-locomotion, COC and AMPH produce these effects through different mechanisms. COC competitively inhibits DAT by binding to a site overlapping with that of DA (Beuming et al., 2008; Chen et al., 1998; Huang et al., 2009; Sitte et al., 1998; Sonders et al., 1997), whereas AMPH has a more complex interaction with the DA system (Sitte and Freissmuth, 2015; Sulzer et al., 2005). AMPH is a DAT substrate that, not only competitively inhibits DAT, but also increases cytoplasmic DA levels both by reducing DA packaging into synaptic vesicles (Freyberg et al., 2016) and increasing DA synthesis (Larsen et al., 2002), the latter leading to non-exocytic DA release via reverse transport through DAT (Sitte and Freissmuth, 2015). Although DA efflux has been established to occur in rodent brain slices by several groups (Jones et al., 1998; Schmitz et al., 2001), as well as *Drosophila* (Freyberg et al., 2016), recent studies suggest that this mechanism of action might not play a dominant role in freely moving mice and rats (Daberkow et al., 2013; Siciliano et al., 2014).

Male mice habituated to the open field maze were administered saline or AMPH (2.5 or 5 mg/kg, i.p.) and monitored for locomotion. Similar to Flot1^fl/fl^ controls, Flot1 cHet demonstrated the predictable, hyperlocomotor response to both doses of AMPH (Figure 2A). In contrast, Flot1 cKO mice demonstrated a significantly blunted response, showing little to no detectable locomotor response at 2.5 mg/kg and a significantly reduced response at 5 mg/kg (Figure 2A). This diminished response was specific to AMPH, because the ability of Flot1 cKO mice to respond to COC was indistinguishable from controls (Figure 2B). Examination of basal locomotion also revealed that the ability of Flot1 cKO mice to habituate to the open field was unaffected by loss of Flot1 expression (Figure 2C). Similar results were obtained with Flot1 cKO female mice (Figure S2A). These data demonstrate that loss of Flot1 in DAergic neurons can separate the ability of mice to respond to AMPH *versus* COC *in vivo*, indicating that in mice AMPH exerts effects beyond DAT inhibition alone.

**Figure 2.**
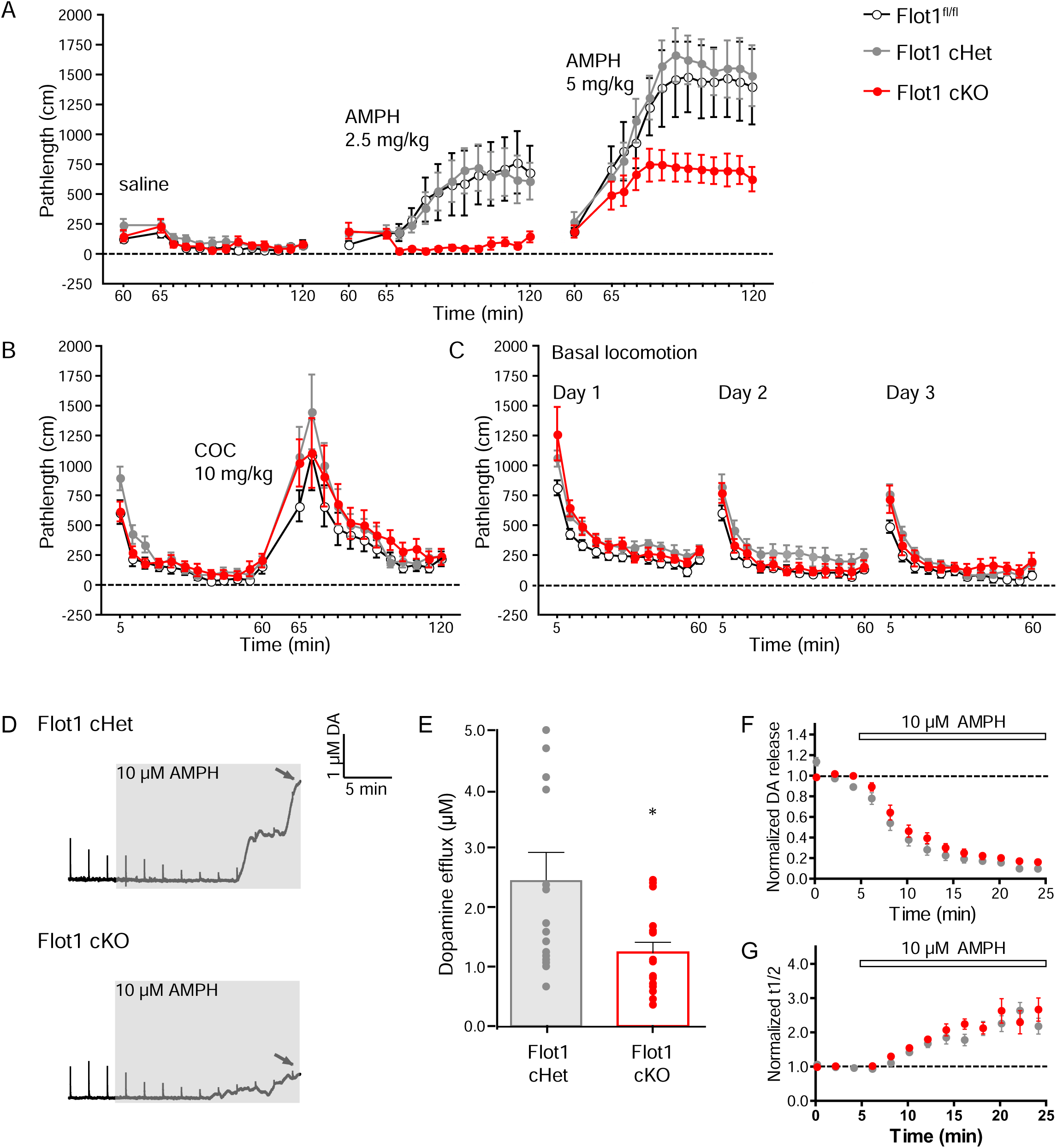
Conditional elimination of Flot1 in DA neurons impairs the ability of mice to respond to AMPH *in vivo*. **A.** Flot1 cKO mice show diminished response to AMPH, but normal response to COC. Flot1^fl/fl^ (black), Flot1^fl/+^::DAT^iresCre+/-^ (Flot1 cHet, gray) and Flot1^fl/fl^::DAT^iresCre+/-^ (Flot1 cKO, red) littermate male mice were habituated in an open field maze for 60 min, then administered saline, or 2.5 or 5 mg/kg of amphetamine (AMPH) i.p. and monitored for an additional 60 min. Flot1^fl/fl^ and cHet mice demonstrate a typical, hyperlocomotor response to 2.5 and 5 mg/kg AMPH. In contrast, cKO mice show no response at 2.5 mg/kg and a significantly diminished response at 5 mg/kg. (Repeated measures (RM)-ANOVA was used to determine the effect of genotype and treatment (Rx) on Pathlength: ***Saline:*** Genotype effect (F_(2,34)_=1.299, p=0.2849). ***AMPH 2.5***: Genotype effect (F_(2,34)_=3.726, p=0.0365). Rx effect (F_(2,12)_=8.896, p<0.0001). Interaction in Genotype and Rx (F_(2,24)_ = 3.461, p <0.0001). Fisher PLSD reveals a difference in Flot1^fl/fl^ and cKO (p=0.0215), cHet and cKO (p=0.0323) but not Flot1^fl/fl^ and cHet (p=0.8623). ***AMPH 5***: Genotype effect (F_(2,34)_=5.005, p=0.0124). Rx effect (F_(2,12)_=25.025, p<0.0001). Interaction in Genotype and Rx (F_(2,24)_ = 3.190, p <0.0001). Fisher PLSD reveals a difference in Flot1^fl/fl^ and cKO (p=0.0213), cHet and cKO (p=0.0057) but not Flot1^fl/fl^ and cHet (p=0.6020)). **B.** Locomotor response to 10 mg/kg of cocaine (COC) i.p. The hyperlocomotor response was indistinguishable across genotype. (RM-ANOVA finds no effect of genotype on Pathlength (F_(2,34)_=1.679, p=0.2016.)) **C.** Basal locomotion and habituation to the open field was indistinguishable across genotype. (RM-ANOVA finds no effect of genotype on Pathlength on any day (Day 1: Genotype effect (F_(2,34)_ = 2.108, p = 0.1371); Day 2: Genotype effect (F_(2,34)_ = 3.241, p = 0.0675); Day 3: Genotype effect (F_(2,34)_ = 0.324, p = 0.7252)). For A-C, analysis of female mice can be found in Figure S2. n = 12, 12, 13 (Flot1^fl/fl^, cHet, cKO). Data represented as mean ±S.E. **D-G.** Cyclic voltammetry in slice preparation from Flot1 cHet and cKO mice. Flot1 cHet mice were used given that they were indistinguishable in behavioral tasks from Flot1^fl/fl^ mice, but control for the presence of DAT^iresCre^, which has previously been shown to diminish endogenous DAT levels. **D.** Representative traces from slice recordings from cHet (top) or cKO (bottom). The peak of the spikes represents electrically stimulated vesicular DA release, or evoked release. Each peak, recorded every 2 min, is a single-pulse electrical stimulation. The arrow indicates the time when data was taken for the change in baseline, which represents AMPH-induced efflux. This was approximately 20 min following the perfusion of 10 μM AMPH onto slices. **E.** Quantification of DA efflux revealed a significantly reduced response in Flot1 cKO slices (Mann Whitney U test reveals a significant difference between cHet and cKO mice: p = 0.0381. N = 5 mice, n = 15 slices/genotype). **F, G.** Quantification of DA release (**F**) and reuptake represented as t_1/2_ (**G**) after 10 µM AMPH. AMPH inhibited both DA release and DA reuptake similarly across genotypes. Similar findings were made using chronoamperometry in slice (Figure S2B). Data represented as mean ± S.E. histogram with individual data points shown.

Reminiscent of these behavioral findings, we previously reported that depletion of Flot1 in primary mesencephalic cultures of DA neurons using RNA interference leads to diminished DA release in response to AMPH (Cremona et al., 2011). To determine if we observe similar changes in these mice, we used cyclic voltammetry to characterize the release and uptake dynamics of DA in response to AMPH using striatal slices from littermate Flot1 cHet and cKO mice (Figures 2D-G). cHet mice were used to control for the presence of DAT^iresCre^ (Backman et al., 2006). Quantification revealed that whereas AMPH-mediated inhibition of DA release and reuptake were similar across genotypes, non-exocytic release of DA in response to AMPH was significantly reduced in cKO mice (Figures 2D and 2E). Similarly, chronoamperometry studies in slice demonstrated that release of DA in response to AMPH was blunted in cKO mice (Figure S2B). In contrast, no significant differences were observed in the amplitude and t_1/2_ of stimulation-dependent DA release for AMPH-induced changes (Figures 2F and 2G), indicating that vesicular DA depletion and inhibition of DAT are the same between the genotypes. Taken together, these results suggest that Flot1 is required selectively for the ability of DAT to reverse-transport DA in response to AMPH without affecting DAT-mediated DA uptake.

### The membrane milieu of DAT is important for the ability of DAT to respond to AMPH

By performing a spatially restricted KO of Flot1, we have revealed that AMPH-induced events and the membrane partitioning of its substrate DAT are significantly diminished. Previous studies using a global, constitutive deletion of *Flot1* however have reported only modest phenotypic changes (Bitsikas et al., 2014; Bodrikov et al., 2017; Ludwig et al., 2010). This was surprising given the importance attributed to Flotillin-mediated partitioning of cell adhesion molecules (CAMs) such as E-cadherin. A recent study using RNA-based rather than genomic approaches in order to deplete expression of Flotillins revealed that loss of the Flot1/Flot2 hetero-oligomer disrupts vertebrate embryogenesis via CAM scaffolding (Morris et al., 2017), suggesting that the developmentally-driven, epigenetic changes that ensue following genomic modifications (El-Brolosy and Stainier, 2017; Rossi et al., 2015) may mask Flot1-function. We hypothesize that, although Flot1 has been implicated in various cell-based processes, the function of Flot1 is to scaffold substrates such as CAMs or DAT to cholesterol-rich membranes. To test this hypothesis, we used Cre-based approaches to create two additional lines of mice: A traditional knockout model of Flot1 (Flot1 KO) that deletes *Flot1* at early stages of development (Figure 3A-E); and an inducible KO model of Flot1 (Flot1 iKO) that enables deletion of *Flot1* in adult mice (Figures 3F-J). Using these mice, we determined how eliminating Flot1 at different time points might affect both Flot1-mediated DAT partitioning and Flot1-dependent AMPH-induced behaviors.

**Figure 3.**
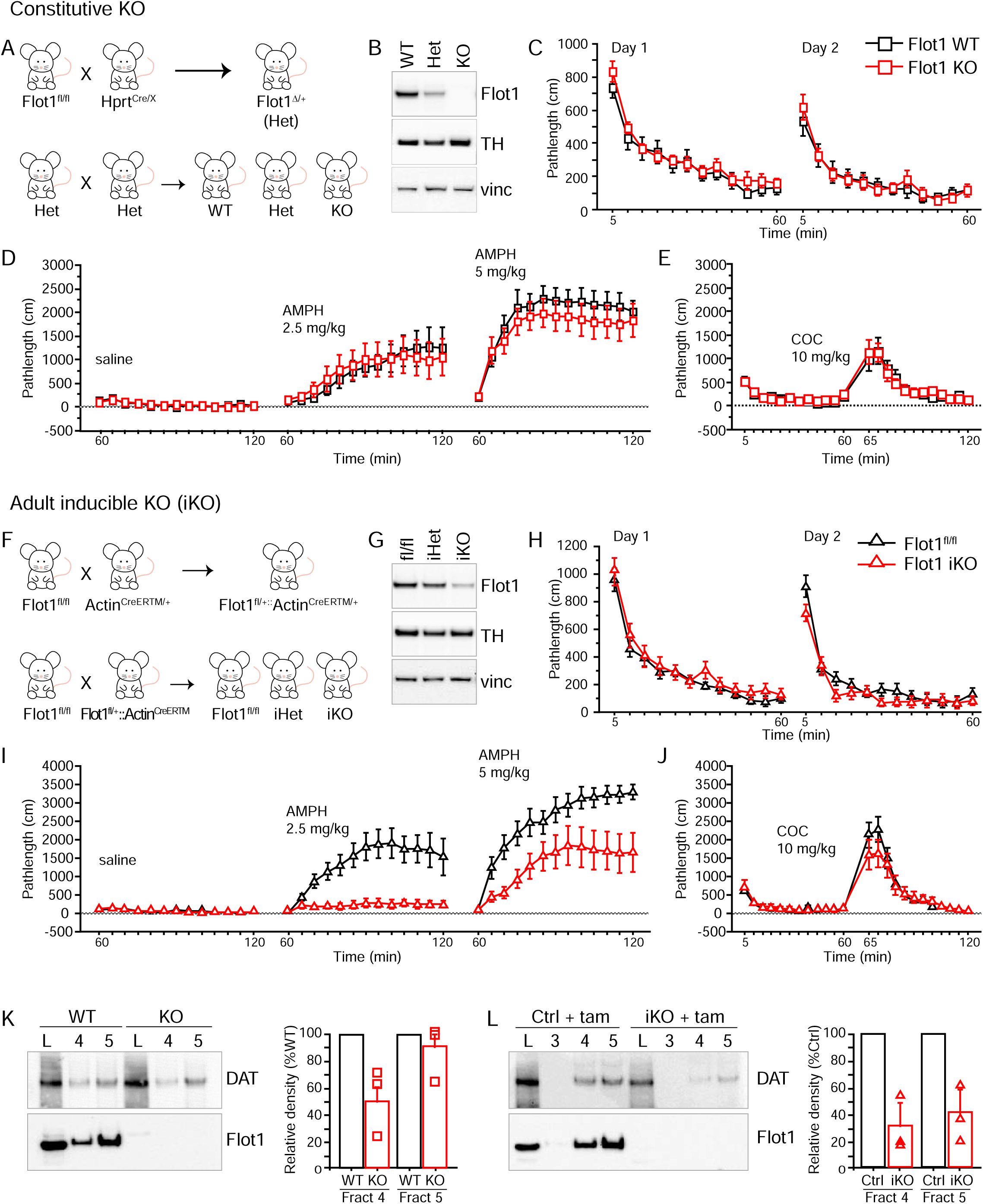
Developmental events mask the ability of Flot1 to scaffold DAT into DRMs as revealed by constitutive versus conditional genetic approaches. **A-E.** Constitutive loss of Flot1 does not demonstrate a blunted response to AMPH. **A.** Breeding schema for the creation of the heritable *Flot1*-deleted (Δ) allele and constitutive *Flot1* knockout mice (Flot1 KO). Flot1 heterozygous (Het) mice are intercrossed to create the littermates used in the experiments. **B.** Immunoblotting of substantia nigra lysates WT, Het and KO littermates for Flot1. Blots were also probed for TH to determine relative levels of DA. Vinculin (vinc) serves as a loading control. Similar results were obtained for the whole brain (Figure S3). **C-E.** Locomotor behavior of mice in the presence or absence of AMPH or COC as performed in Figure 2. In contrast to cKO mice, KO mice show a normal response to AMPH. (RM-ANOVA was used to determine the effect of genotype and Rx on Pathlength: ***Basal***: Day 1: Genotype effect (F_(2,30)_=1.031, p=0.3689); Day 2: Genotype effect (F_(2,30)_=0.644, p=0.5321). ***Saline:*** Genotype effect (F_(2,30)_=0.257, p=0.7752). ***AMPH 2.5***: Genotype effect (F_(2,30)_=0.429, p=0.6553). Rx effect (F_(12,360)_=14.985.896, p<0.0001). ***AMPH 5***: Genotype effect (F_(2,30)_=0.189, p=0.8286). Rx effect (F_(12,360)_=32.954, p<0.0001). ***COC 10***: Genotype effect (F_(2,30)_=.393, p=0.6783)). n=12/genotype. Data represented as mean ±S.E. **F-J**. Conditional adult-inducible deletion of Flot1 recaptures the Flot1-dependent response to AMPH. **F.** Breeding schema for the creation of the Flot1 inducible KO mouse (Flot1 iKO). Crossing the conditional Flot1 allele with a tamoxifen-inducible Cre line, Actin^CreERTM^ ultimately leads to the creation of mice in which Flot1 can be deleted in a temporally regulated manner through the administration of tamoxifen (tam). **G.** Administration of tam at 4 wks of age leads to efficient loss of Flot1. Immunoblotting of substantia nigra lysates 4 weeks after the last injection of tam. Whole brain lysates are shown in Figure S3. **H-J.** Open Field paradigm performed in 9 week-old mice. Unlike the Flot1 KO mice, but similarly to the Flot1 cKO mice, tam-treated Flot1 iKO mice showed a diminished response to AMPH, whereas basal locomotion and COC-induced hyperlocomotion was normal. (RM-ANOVA was used to determine the effect of genotype and Rx on Pathlength: ***Basal***: Day 1: Genotype effect (F_(2,27)_=2.388, p=0.1109); Day 2: Genotype effect (F_(2,27)_=1.863, p=1.746). ***Saline:*** Genotype effect (F_(2,27)_=1.095, p=0.3488). ***AMPH 2.5***: Genotype effect (F_(2,27)_=11.171, p=0.0003). Rx effect (F_(12,324)_=13.792, p<0.0001). Interaction in Genotype and Rx (F_(24,324)_ = 5.550, p <0.0001). Fisher PLSD reveals a difference in Flot1^fl/fl^ and iKO (p=0.0002). ***AMPH 5***: Genotype effect (F_(2,27)_=4.839, p=0.0160). Rx effect (F_(12,324)_=38.337, p<0.0001). Interaction in Genotype and Rx (F_(24,324)_ = 2.415, p =0.0003). Fisher PLSD reveals a difference in Flot1^fl/fl^ and iKO (p=0.0174). ***COC 10***: Genotype effect (F_(2,27)_=0.257, p=0.7754). n=10/genotype. Data represented as mean ±S.E. Similar results were obtained in Flot1 iKO female mice (Figure S3). **K, L.** DAT from Flot1 iKO but not Flot1 KO mice fails to partition into DRMs. Sucrose density gradients (SDG) centrifugation on 1% Brij58 striatal lysates from Flot1 KO (K) and iKO (L) mice. To quantify the relative partitioning of the genetically modified mice to their respective controls, the Flot1-positive DRM fractions and total lysates of control and experimental genotypes were run together for immunoblotting. Fractions were normalized to total lysate. Quantification revealed that in Flot1 KO lysates, a small change of DAT partitioning in the more buoyant fraction 4 was observed, whereas in Flot1 iKO lysates, a significant decrease in DAT partitioning into both the DRM fractions was observed. Mann Whitney U test of Ctrl (WT) vs. Flot1 KO fractions reveals a significant difference in fraction 4 (p = 0.045) but not in fraction 5 (p = 0.4867). Similar test of Ctrl (Flot1^fl/fl^) + tam vs. Flot1 iKO + tam reveals a significant difference in both fractions 4 (p = 0.045) and 5 (p = 0.045). n = 3/genotype. Data represented as mean ± S.E., with individual data points also shown.

To create Flot1 KO mice, Flot1^fl/fl^ mice were crossed with *Hprt*^Cre/+^, which deletes *Flot1* in the oocyte, thereby giving rise to a heritable *Flot1*-deleted (Δ) allele (Figures 3A, B and S3A, B). The open field paradigm revealed that global, constitutive loss of Flot1 has no impact on basal locomotion (Figure 3C), and, consistent with previous reports (Bitsikas et al., 2014; Bodrikov et al., 2017; Ludwig et al., 2010), has no obvious phenotype. We next examined the locomotor response of Flot1 KO and littermate WT mice to AMPH and COC (Figure 3D and 3E). In contrast to the conditional elimination of *Flot1* in DAergic neurons alone, global Flot1 KO has no effect on either AMPH- or COC-induced locomotion compared to controls.

Next, Flot1^fl/fl^ mice were crossed with Actin^CreERTM /+^ mice, which enables Cre/loxP-mediated gene deletion upon administration of tamoxifen (tam), to permit global deletion of *Flot1* in adult mice (Figures 3F, G and S3C, D). Western blots of SN and striatum from Flot1 iKO mice revealed significant deletion of *Flot1* 4 weeks post-injection (Figures 3G and S3D). Tam-injected Flot1 iKO and Flot1^fl/fl^ littermate controls were assessed for a locomotor response to AMPH and COC (Figures 3H-J). No differences in basal locomotion and habituation to the open field (Figure 3H) nor COC-induced hyperlocomotion were observed across genotypes (Figures 3H-J). In contrast to the Flot1 KO mice however, tam-injected iKO mice demonstrated a blunted response to AMPH, similar to that observed in Flot1 cKO mice; no response was observed at 2.5 mg/kg and a significantly diminished response at 5 mg/kg (Figure 3I). The female cohort of mice behaved similarly (Figure S3E). These data strongly support the model that global, genomic deletion of *Flot1* throughout development leads to changes that mask the impact of Flot1 deletion.

We next determined whether the changes in behavioral outcome between Flot1 KO and tam-injected Flot1 iKO strains are reflected in the partitioning of DAT. Sucrose density gradient centrifugation was performed on striatal lysates generated from Flot1 KO and tam-injected Flot1 iKO mice and their respective controls. The Flot1-positive DRM fractions and total lysates from control and experimental genotypes were run together on Western blots to permit quantification with respect to controls (Figures 3K and 3L). The degree of DAT partitioning correlated with the behavioral response to AMPH. In the Flot1 KO mice which showed a normal AMPH-induced response, only a small change in DAT partitioning was observed in striatal gradient (Figure 3K), as described for amyloid Aβ peptide (Bitsikas et al., 2014). In contrast, loss of Flot1 in the Flot1 iKO mice led to a significant decrease of DAT partitioning into all DRM fractions (Figure 3L), reminiscent of our observations in Flot1 cKO mice (Figure 2A). Thus, lysates from mice whose AMPH-induced response is diminished by the loss of Flot1 (Flot1 cKO and Flot1 iKO) also demonstrate diminished partitioning of DAT into DRMs. Moreover, lysates from mice that maintain their ability to respond to AMPH despite the loss of Flot1 (Flot1 KO) also maintain DAT partitioning. These data further strengthen the correlation between the ability of DAT to reverse-transport DA in response to AMPH and the ability of DAT to partition into cholesterol-rich DRMs.

### Loss of Flot1 specifically disrupts partitioning of a subset of proteins

As we have shown above, conditional loss of Flot1 excludes DAT from the 1% Brij58 DRMs. A simplistic interpretation of these observations is that loss of Flot1 generally disrupts partitioning of proteins throughout the membrane. To explore this further, we examined partitioning of various neuronal proteins that have been previously shown to occupy cholesterol-rich membranes (Figure 4A). We found the loss of Flot1 does not disrupt the distribution of proteins such as SNAP-25 or Src. Partitioning of the paralogue Flot2 on the other hand was notably disrupted, suggesting that hetero-oligomerization of Flot1 and Flot2 might be critical for their mutual function as suggested previously (Ludwig et al., 2010). Taken together, although the conditional deletion of *Flot1* disrupts partitioning of DAT, this does not indicate global alterations in membrane protein partitioning upon loss of Flot1.

**Figure 4.**
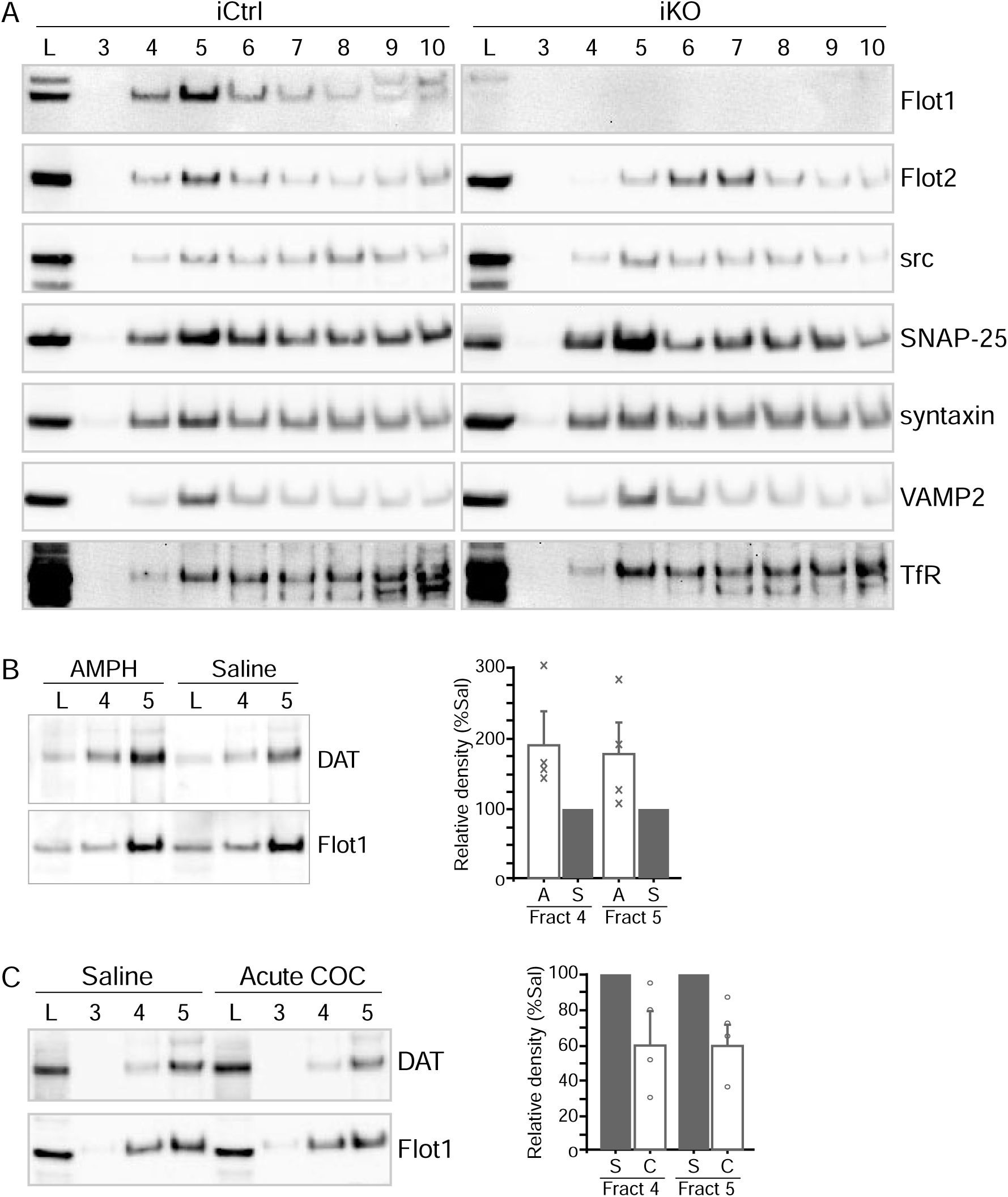
DAT activity modulates the partitioning of DAT in cholesterol-rich membranes. **A. Sucrose density gradients reveal that Flot1 is not required to scaffold all proteins into DRMs.** Striatal lysates from Flot1 iCtrl and iKO mice were fractionated across a discontinuous sucrose gradient (SDG) as described above. Fractions were collected from top (2) to bottom (10) of the gradient. A panel of DRM-associated proteins did not shift with the loss of Flot1, except Flot2, whose DRM-association has previously been shown to be dependent on Flot1. **B.** AMPH increases the detectable levels of DAT in DRMs. Striatal homogenates from WT mice treated with a single injection of 5 mg/kg (i.p.) of AMPH or saline were lysed in 1% Brij58, and run on SDGs. AMPH treatment led to a significant increase in the amount of DAT in the Flot1-positive DRM fraction (Mann-Whitney U: Fraction 4, p=0.0209, Fraction 5, p=0.0209; n=4). AMPH treatment had no detectable impact on the distribution of DAT in Flot1(DAT) cKO striata (Fig. S4A). **C.** Inhibition of DAT via COC diminishes the detectable levels of DAT in DRMs. Striatal homogenates from WT mice treated with a single injection of 20 mg/kg (i.p.) of COC or saline were lysed in 1% Brij58, and run on SDGs. In contrast to AMPH, COC leads to a significant decrease in the amount of DAT in the DRM fractions (Mann-Whitney U: Fraction 4, p=0.0209, Fraction 5, p=0.0209; n=4).

### AMPH and COC differentially influence partitioning of DAT into detergent-resistant, cholesterol-rich membranes

We next performed a series of studies to determine why DAT partitioning is required for AMPH-induced events. Recent studies suggest that, similarly to Flot1, DAT is subject to palmitoylation, a reversible post-translational modification that increases the propensity of proteins to occupy sterol and sphingolipid membranes (Foster and Vaughan, 2011; Moritz et al., 2015). Given that increased palmitoylation of DAT is associated with increased transport rates, and AMPH uptake by DAT should be structurally similar to that of DA (Cheng et al., 2015), we hypothesized that AMPH administration would be sufficient to increase DAT partitioning into cholesterol-rich membranes. DRMs derived from striatal homogenates of WT mice treated with a single injection of 5 mg/kg (i.p.) of AMPH or saline were probed for DAT and Flot1. AMPH treatment led to a significant increase in the amount of DAT in the Flot1-positive DRM fraction, relative to saline treatment. This increase was Flot1-dependent, as AMPH treatment had no detectable impact on the distribution of DAT in Flot1 cKO striata (Figure S4A). To complement this experiment, we next inhibited DAT function with COC (Figure 4C). DAT inhibition by a single i.p. administration of 20 mg/kg COC was sufficient to evoke a measurable decrease in DAT partitioning to DRMs (Figure 4C). This is consistent with the behavioral data from the Flot1 cKO and Flot1 iKO mice, which indicate no relationship between DAT partitioning and COC-evoked events. In sum, these data show that DAT activity leads to increased DAT partitioning in a Flot1-dependent manner, and inhibition of DAT activity diminishes this partitioning event.

### Membrane partitioning of DAT is not essential for its phosphorylation

One hypothesis often put forth for membrane microdomain function is that these protein-lipid structures serve to increase efficiency of post-translational modifications to target proteins (Levental et al., 2010). In the case of DAT, phosphorylation of its NH3-terminus, such as at residues Ser7 and Ser12 by protein kinase C and CaMKII α/β, has been shown to be necessary for DA efflux in response to AMPH, (Foster et al., 2002; Khoshbouei et al., 2004; Pizzo et al., 2014). We therefore examined whether phosphorylation of DAT at these residues requires Flot1 (Figures S4). Because the antibody against phospho-Ser7 is specific for human DAT (Karam et al., 2017), we turned to a well-established heterologous expression system for DAT, namely YFP-DAT stably expressed in modified HEK293 cells (EM4 YFP-DAT) (Kahlig et al., 2004; Sen et al., 2005). We found that phosphorylated Ser7 can be detected under baseline conditions, which can be readily augmented by PKC activation via PMA as previously reported (Figure S4A) (Cowell et al., 2000; Giambalvo, 1992; Kantor and Gnegy, 1998). Depletion of Flot1 by RNA interference had no detectable impact on basal or PKC-mediated phosphorylation.

To examine phosphorylation of endogenous murine DAT, we also examined Thr53 phosphorylation in DRMs from Flot1 cKO and control striata. Thr53 phosphorylation has been implicated in AMPH-induced efflux, although how and why this phosphorylation event occurs is unclear (Foster et al., 2012). Given that this is the only site at which an antibody against the phosphorylated murine form is available for DAT, we determined the dependence of Thr53 phosphorylation on Flot1 and AMPH (Figure S4F). Immunoblotting of gradient fractions with the phospho-DAT(Thr53) antibody revealed that this residue is constitutively phosphorylated in mouse striata, and such phosphorylation persists despite absence of Flot1 (Figure S4B). Thus, although phosphorylation of the NH_3_-terminal region of DAT has been shown to be a critical regulator of AMPH-induced efflux, our data indicate that the presence of this key modification does not require Flot1 nor DAT partitioning.

### The conformation of DAT differs in cholesterol-rich versus cholesterol-poor membranes

Lipid-protein interactions have been shown to profoundly influence membrane protein structure and function (Dominguez et al., 2016; Dowhan and Bogdanov, 2011; Lee, 2004). For example, the biophysical properties of the membrane microenvironment have been shown to effect the structure and function of ion channels, slowing or accelerating their responsiveness (Tillman and Cascio, 2003). An important role for cholesterol in DAT protein folding has already been proposed based on the structure of dDAT (Penmatsa et al., 2013). We therefore tested if the membrane environment of DAT is important for AMPH-induced reverse transport of DA because this environment is conducive for stabilizing a conformation of DAT that is important for efflux to occur. To examine potential conformational differences of endogenous DAT that might be present in 1% Brij58-resistant membranes, we performed limited proteolysis to determine if different cleavage susceptibilities can be detected across fractions (Figure 5). The fractions were exposed to trypsin, papain, or 2-nitro-5-thiocyanatobenzoic acid (NTCB) for the indicated times, and then processed for Western blot analyses. All three enzymes revealed that the susceptibility of cleavage of DAT notably differed among the fractions: For trypsin and papain (Figures 5A, B), DAT was more susceptible to cleavage in DRMs than in non-DRMs, whereas for NTCB the reverse was true (Figure 5C). BSA controls ensured that differential cleavage was not due to differing sucrose concentrations across fractions (Figure S5). These data indicate that DAT in Flot1-positive DRMs is conformationally distinct from DAT in non-DRM fractions.

**Figure 5.**
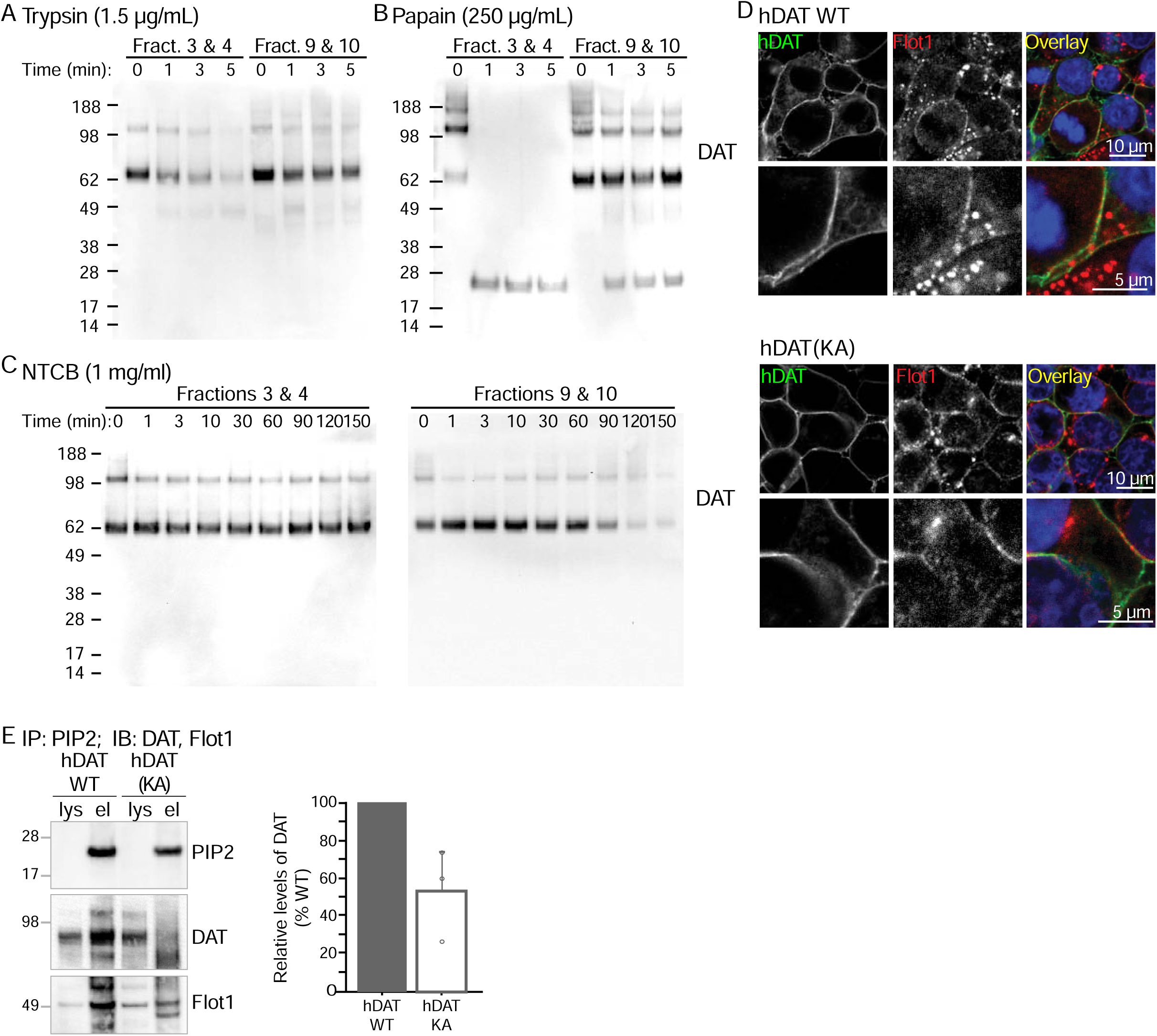
DAT present in 1% Brij58 DRMs is conformationally distinct from non-DRM fractions due to its ability to interact with PIP2. **A.-C.** Limited proteolysis of Flot1-positive *versus* Flot1-negative SDG fractions using trypsin (**A**), papain (**B**) or NTCB (**C**) reveal that the conformation of DAT differs between fractions. The combined Flot1-positive fractions 3 and 4 were compared to the Flot1-negative fractions 9 and 10. **A.** Two different concentrations of trypsin revealed that the Flot1-positive fractions were more susceptible to proteolysis (see also Figure S5). **B.** Papain, which has similar proteolytic cleavage sites to trypsin yielded similar results. **C.** NTCB which has a different proteolytic capacity from trypsin and papain, also yielded differential proteolysis across fractions, but in this case, the Flot1-positive fractions showed increased resistance. BSA controls indicate that the different sucrose levels do not affect proteolytic rates (Figure S5). **D.** Heterologous expression of hDAT or hDAT(K3A, K5A)(hDAT(KA)) in HEK293 cells. hDAT is stained for immunofluorescence with MAB369 (green) and Flot1 with BD#610821 (red). Mutagenesis of the PIP2 binding residues of the N-terminus of DAT to Ala does not grossly affect the membrane localization of DAT. **E.** Co-immunoprecipitation of PIP2 with DAT confirms that DAT Lys3 and Lys5 are important for the ability of DAT to interact with PIP2. 2 to 6 µg of total lysate (Lys) of cells shown in **D** were run against the equivalent of 50 µg of the eluate (el) after immunoprecipitation with an anti-PIP2 antibody. IPs were performed in 1% Tx-100 containing RIPA buffer as described in methods, to ensure disruption of the Brij58 resistant DRMs. Blots were probed for DAT and Flot1. Quantification of the amount of DAT immunoprecipitated confirms that hDAT(KA) is unable to interact as well to PIP2 as hDAT(WT) (right). The eluted amount was corrected for the total level of DAT detected and lysate and the PIP2 immunoprecipitation. Significantly less DAT was retrieved in the hDAT(KA) mutant relative to hDAT(WT). (Mann-Whitney U: p=0.03; n=3)

**Figure 6.**
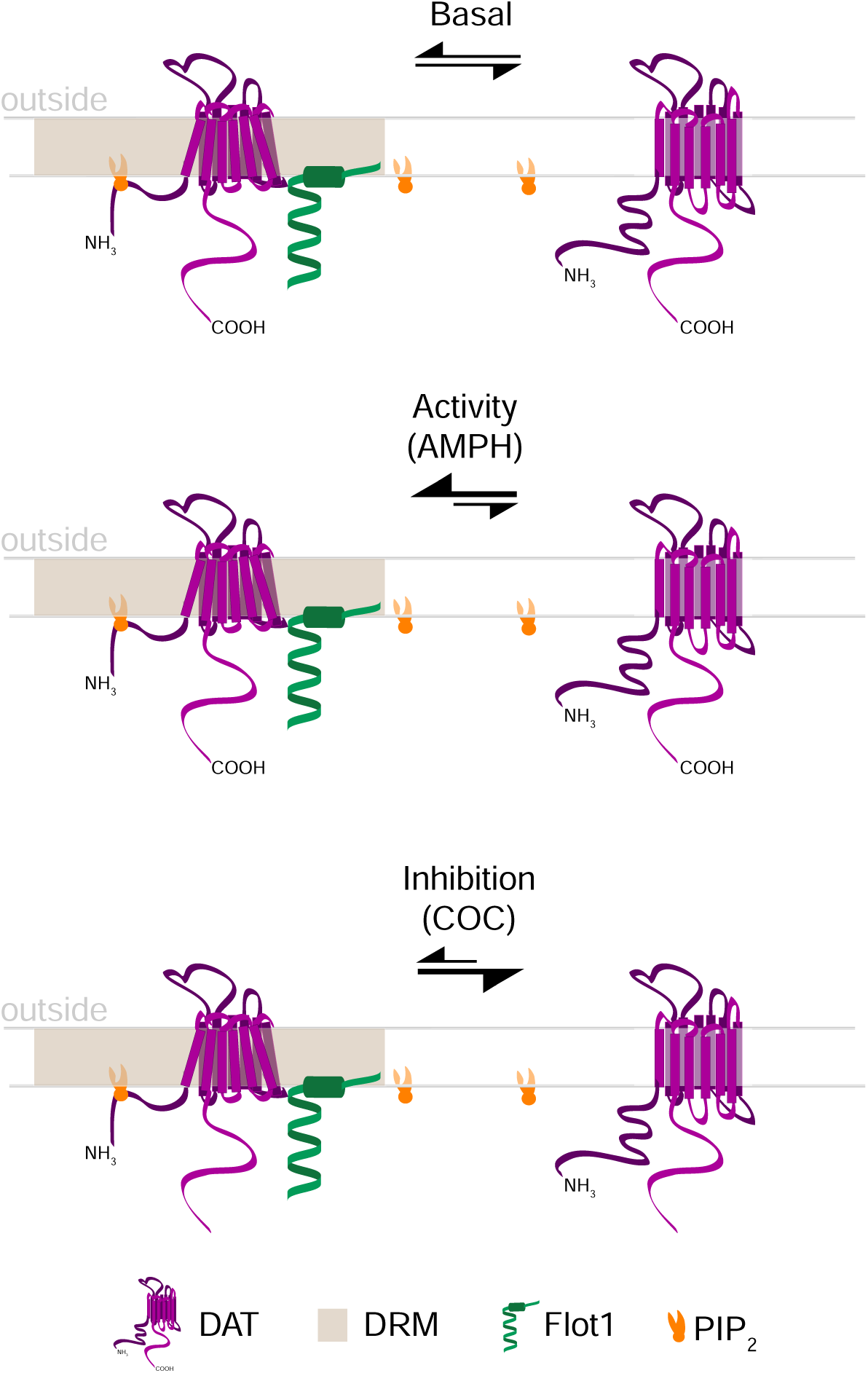
Schematic representation of Flot1-mediated membrane partitioning. Model of Flot1-mediated partitioning of DAT. DAT, like its SLC6 family member SERT, is found in both DRMs and non-DRMs. Administration of AMPH vs. COC can change this partitioning, possible by changes in DAT palmitoylation.

### The conformation of DAT in 1% Brij58 DRMs depends on its N-terminal interaction with PIP_2_

Recently, Lys3 and Lys5 of human DAT were shown to interact with the widely distributed, plasma membrane-enriched phospholipid phosphatidylinositol 4,5-bisphosphate (PIP_2_) and to be necessary for a conformation of DAT that favors DA efflux in the presence of AMPH (Hamilton et al., 2014). Disruption of the PIP_2_ interaction via mutation of heterologously-expressed hDAT diminished AMPH-induced efflux in mammalian cells, as well as the ability of *Drosophila* to mount a hyperlocomotor response to AMPH (Hamilton et al., 2014). We therefore tested whether the conformational difference we detected in the DRMs was due to an interaction between PIP_2_ and DAT. Wild-type hDAT (hDAT(WT)) or hDAT(K3A, K5A) (hDAT(KA)) was transiently expressed into HEK293 cells. Alanine mutagenesis of Lys3 or Lys5 of DAT has no notable impact on the ability of DAT to traffic to the plasma membrane (Figure 5D). Co-immunoprecipitation experiments revealed that even though hDAT(KA) is expressed at the cell surface, its interaction with PIP_2_ is significantly diminished, as previously shown (Figure 5E) (Hamilton et al., 2014). These data suggest that, despite the widespread presence of PIP_2_ in the plasma membrane, the membrane environment of DAT represented by the 1% Brij58-resistant DRMs helps to maintain a conformation consistent with that of DAT interacting with PIP_2_. Although hDAT(KA) was detected in DRMs, we noted that it did not partition as efficiently as hDAT(WT) (Figure S5C), raising the possibility that the interaction between PIP_2_ and DAT might also contribute to maintain DAT in the cholesterol-rich environment of the DRMs. Extending these findings to our results *in vivo*, we propose that in Flot1 cKO and tam-injected iKO mice, the absence of Flot1 prevents DAT from partitioning into sterol-rich membranes, thereby diminishing the ability of DAT to interact with PIP_2_, and its ability to stabilize a conformation that is more effective for AMPH-induced DA efflux. Taken together, our data indicate that the local membrane environment of DAT can modify the stability of key conformations of DAT, and markedly affect its function.

## Discussion

Our initial studies of Flot1-DAT interactions were inspired by the original observation by Haase *et al.* (Magnani et al., 2004), demonstrating not only that the structurally similar SERT partitions into DRMs, but that it did so incompletely. The question of why a transmembrane protein might occupy different membrane environments, as suggested by density gradient centrifugation, led us to hypothesize that the membrane environment is functionally important. Our previous study in cell-based systems revealed that membrane localization of DAT might be relevant for either regulation of DAT cell surface availability, or of AMPH-induced efflux of DA (Cremona et al., 2011). DAT-mediated uptake of DA in EM4 cells has been shown to be Flot1-independent (Cremona et al., 2011). Moreover, studies in *Drosophila* have shown that loss of Flot1 diminished AMPH-induced larval hyperlocomotion, but did not impede the behavioral response to methylphenidate, an inhibitor of DAT uptake (Pizzo et al., 2013). Given that much of the preceding work used heterologous systems or dissociated embryonic cultures, we sought to determine which of these events could be recapitulated in the vertebrate brain.

Using three mouse models that eliminate *Flot1* expression by different means, we were able to distinguish the impact of the membrane environment on the ability of endogenous DAT to respond to psychostimulants at the level of mouse behavior: COC-mediated hyperlocomotion was intact but AMPH-induced efflux and hyperlocomotion were selectively diminished. To our knowledge, our findings represent a unique *in vivo* demonstration of how the local lipid environment of a membrane protein, as implied by DRM isolation, can affect brain function and its ensuing behavior. The loss of Flot1 does not modulate the membrane environment of all proteins, but rather is specific for distinct substrates. Moreover, whereas Flot1 is important for the maintenance of DAT in 1% Brij58-resistant DRMs (Cremona et al., 2011; Hong and Amara, 2010; Jones et al., 2012), substrates such as E-cadherin are maintained in 1% TritonX-100-resistant DRMs (Kurrle et al., 2013), indicating that despite specificity for its substrates, Flot1 associates to membranes of different composition. Thus, the common use of Flot1 as a general membrane raft marker inadvertently obscures the specificity of Flot1 function. Specificity might be achieved in part by lipid composition, but also via interactions between Flot1 and its substrate (Banning et al., 2011; Cremona et al., 2011; Kurrle et al., 2013). It is likely that the structurally similar Flot2 also contributes significantly to this specificity. Although we have not examined Flot2, several studies have established a tight inter-relationship between these two proteins (Banning et al., 2011; Ludwig et al., 2010; Solis et al., 2007). For example, Flot2 does not partition into DRMs when Flot1 is absent (Ludwig et al., 2010), and the protein stability of Flot1 depends on the presence of Flot2 (Solis et al., 2007). We also find that endogenous DAT, Flot1 and Flot2 co-immunoprecipitate in striatal lysates (data not shown). How the interaction of Flotillins with their substrates occur is unclear, but given that DAT activity can alter its partitioning, events such as post-translational modifications of substrates such as DAT or the Flotillins themselves might be key to complex formation (Cremona et al., 2011).

The necessity for partitioning of DAT into cholesterol-rich membranes for AMPH-related events gave rise to different possibilities. First, it is notable that disrupting partitioning did not impact reuptake by DAT, indicating that the partitioning we observed was independent of the known cholesterol association with DAT to maintain protein folding and structure. Our cyclic voltammetry studies indicate that the cell surface availability of DAT under basal conditions or in the presence of AMPH is not grossly affected upon conditional loss of Flot1, thus we focused on DA efflux. Previous studies have revealed post-translational modifications of DAT, namely phosphorylation (Foster et al., 2012; Khoshbouei et al., 2004), as well as various protein and lipid interactions with DAT (Eriksen et al., 2010; Garcia-Olivares et al., 2017; Hamilton et al., 2014; Khelashvili et al., 2015a; Khelashvili et al., 2015b), are important for DAT efflux. Although membrane microdomains are often thought to act as platforms for post-translational events (Simons and Ikonen, 1997), we surprisingly found that DRM localization is not required for phosphorylation of DAT, neither on Ser7 in heterologous hDAT (Karam et al., 2017) nor Thr53 in endogenous mDAT (Foster et al., 2012). Heterologous experiments in cells suggest that nystatin can diminish phosphorylation of DAT Ser7, and chronic phosphorylation can rescue amphetamine activity despite the absence of of Flot1 in *Drosophila* (Karam et al., 2017). Although the partitioning phenotype of hDAT in fly neurons remains unknown, one potential interpretation given our findings is that, rather than promoting phosphorylation, partitioning of DAT might protect Ser7 from dephosphorylation, thereby increasing steady state levels of phosphorylated DAT. Better understanding of the phosphorylation status of endogenous DAT under basal or drug-treated conditions is necessary to extend these studies.

Computational and functional experiments demonstrate that the interaction of PIP_2_ with the NH_3_-terminus of DAT promotes a conformational transition of DAT that is relevant to AMPH-induced efflux (Hamilton et al., 2014; Khelashvili et al., 2015a). PIP_2_ is found throughout the membrane (van Rheenen et al., 2005), and thus the basis for DAT-PIP_2_ enrichment in DRMs is unclear. One possibility is that the stabilizing effects of cholesterol-rich membranes is relevant; previous structure-function studies suggest that cholesterol enrichment in plasma membranes leads to increased protein activity, and that lipid-protein interactions are necessary for proteins to achieve optimal function (Semrau and Schmidt, 2009). In addition, models such as the flexible surface model (FSM), which posits that the membrane is highly elastic, suggest that conformational changes either in response to protein movement or to help facilitate movement, via curvature sensing (Brown, 1994; Gibson and Brown, 1991, 1993), may also explain what happens with DAT when it effluxes. This also implies that the protein can alter the bilayer curvature to facilitate its structural changes. Therefore, DAT may require and cooperate with cholesterol and PIP_2_ in the DRMs to bend the bilayer, which in turn tugs at DAT to drive it to consistently reverse-transport DA.

Although we find that conditional loss of Flot1 has a significant and specific impact on AMPH-induced hyperlocomotion and DA efflux, this Flot1-dependence is only partial; although the response was abolished at 2.5 mg/kg (i.p.) AMPH, it was only blunted at 5 mg/kg. Cyclic voltammetry and amperometry experiments also revealed a significant but partial inhibition at saturating doses of 10 µM AMPH. First, as suggested by experiments using DAergic neurons and flies (Cremona et al., 2011; Pizzo et al., 2013), specific loss of AMPH-evoked responses supports the notion that DA efflux significantly contributes to AMPH-induced hyperlocomotion even at low doses. This is consistent with recently published work indicating a requirement for acute VMAT function at low doses of methamphetamine (Freyberg et al., 2016). What remains unclear, however, is why efflux can still occur at higher doses. Although Flot1-independent competitive inhibition of DAT by AMPH at higher doses might contribute more significantly, it is possible that after significant stimulation, the need for Flot1 to partition DAT might be overcome. For example, AMPH might increase DAT palmitoylation sufficiently to push DAT to partition even in the absence of Flot1, or might causes lipid modifications that change their fluidity. That DAT partitions in constitutive Flot1 KO mice strongly indicates that alternative means can contribute. This question might be resolved in future studies using chronic AMPH paradigms. For example, chronic AMPH treatment may overcome loss of partitioning in Flot1 cKO or iKO mice. Should chronic AMPH prove to be insufficient to drive partitioning and hence efflux, an increased susceptibility to AMPH cytotoxicity in the absence of Flot1 might emerge.

Employing different complementary genetic approaches to eliminate *Flot1* expression was essential for this study, highlighting the well-known adaptability of mice to constitutive, genetic modifications. These data likely reflect the importance of Flot1 in the developing organism, such as the role Flot1 plays in clustering cadherins (Kurrle et al., 2013). Cadherin function is fundamental to embryonic development, yet similar to the previously published Flotillin KO mice, our constitutive Flot1 KO mice are viable (Banning et al., 2011; Bitsikas et al., 2014; Ludwig et al., 2010). In contrast, RNA-mediated depletion of Flotillin expression in zebrafish caused failure at the earliest stages of embryogenesis, an outcome more consistent with what is known about E-cadherin function. The concept that epigenetic changes might lead to developmental compensations is often raised to diminish apparently incongruent or disparate observations made in constitutive KO mice. A recent study indicates how changes at the level of the genome rather than mRNA can lead to discrete transcriptional responses that protect the organism (Rossi et al., 2015). Nonetheless, how genetic loss of Flot1 is masked in the Flot1 KO mice is an interesting question, and will likely lead to further insight into the importance of membrane-lipid interactions in physiologic function. Given that we have seen no changes in expression of other potential DRM-scaffolding proteins by RNAseq or a limited immunoblotting screen (data not shown), and that membrane lipids might normally be maintained at an equilibrium that is close to domain formation, alterations in lipid composition or arrangement within the plasma membrane might be sufficient to drive partitioning of proteins such as DAT into DRMs, in the absence of a catalysts such as Flot1. Nevertheless, these studies indicate that inducible approaches to re-examine the role of Flot1 and other DRM-associated proteins, such as the caveolins, might provide new insights that constitutive-based approaches have over-looked.

Over time the study of membrane biology has evolved from studying the discrete contributions of lipids and membrane proteins to an appreciation of how these two critical components act together to drive protein function. These cooperative events might be driven by lipid composition, which affects membrane architecture, through which subtle yet active restructuring of the membrane serves to modulate a large array of protein activity. Our findings have expanded our understanding of a role for Flot1 in the CNS and used Flot1 as a genetic tool to establish physiological relevance to how membranes influence protein function in the brain. Through these genetic approaches, we also highlight how potential epigenetic changes may have masked the wide array of functions to which proteins such as Flot1 might be relevant.

## Author Contributions

The experimental design was conceived by W.M.F., K.E., S.J.C, E.V.M., J.A.J. & A.Y.; and carried out by, W.M.F., K.E., S.J.C, I.R., C.W.J., & A.Y. The manuscript was written by W.M.F. & A.Y., with contributions from, C.W.J., K.E., E.V.M., & J.A.J. Experiments were supervised by K.E., E.M. & A.Y.

## Acknowledgements

The authors would like to thank Drs. Aurelio Galli, David Sulzer, Tom Melia, Liz Miller, Margaret Rice, Dritan Agalliu and Tyler Cutforth for critical input and discussions, and Mr. Kelvin Pau for initial insight into the work. These studies were supported by The Parkinson Disease Foundation (WMF, CWJ, AY), NIDA 2P01DA012408 (WMF, JAJ, AG, AY), NINDS NS075222 (EVM), and 1R01DA041510 (JAJ).

## Methods

### Antibodies

The following antibodies were used for immunoprecipitation and/or western blot: rat anti-DAT (1:500; EMD Millipore MAB369), mouse anti-Flot1 (1:250; BD #610821), rabbit anti-Flot1 (1:250; Abcam ab41927), rabbit anti-Vinculin (1:5000; Invitrogen 700062), rabbit anti-tyrosine hydroxylase (1:1000; Calbiochem #657012), rabbit anti-CaMKII (1:1000; Cell Signaling Technology #3362), rabbit anti-phosphoDAT (pSer7) (Karam et al., 2017), rabbit anti-phosphoDAT (pThr53) (1:1000; PhosphoSolutions p435-53), mouse anti-PIP_2_ (1:500; Enzo ADI-915-052-020), mouse anti-γ-tubulin (1:10,000; Abcam ab11316), rabbit anti-Src (1:250; Cell Signaling Technology #2108S), mouse anti-SNAP25 (1:2000; ascites), mouse anti-syntaxin (1:1000; ascites), rabbit anti-VAMP2 (1:1000; EMD Millipore AB5856), mouse anti-Transferrin receptor (1:500; Invitrogen 136800). Rat anti-dopamine transporter (1:1000; EMD Millipore MAB369) and rabbit anti-tyrosine hydroxylase (1:2000; Calbiochem #657012) were used for immunohistochemistry. Rat anti-DAT (1:250; EMD Millipore MAB369) and mouse anti-Flot1 (1:100; BD #610821) were used for immunofluorescence.

### Plasmids

pCI-Hygro vectors expressing hDAT(WT) and hDAT(KA) (Lys3 and Lys5 both mutated to Ala were made as previously described (Hamilton et al., 2014).

### Transfections

siRNAs (first duplex sequence: 5’-CUU CAG UUC GUA AUC UCU CUG UGC CUU-3’ and 5’-GGC ACA GAG AGA UUA CGA ACU GAA G-3’; second duplex sequence: UUA UAG AUC UCC UCC ACA GUC AUG UGC-3’ and 5’-ACA UGA CUG UGG AGG AGA UCU AUA A-3’) (Integrated DNA Technology) were used as a mix against Flot1 using Lipofectamine 2000 (Life Technologies) according to manufacturer’s instructions. hDAT(WT) and hDAT(KA) were transfected using Lipofectamine 2000 (Life Technologies) according to manufacturer’s instructions.

### Cell culture

HEK293T cell cultures were maintained in high-glucose Dulbecco’s modified eagle medium (DMEM; Life Technologies) supplemented with 10% fetal bovine serum (FBS; Life Technologies) at 37°C and in a 5% CO_2_-containing atmosphere.

EM4-YFP-DAT were created and maintained as previously described (Kahlig et al., 2004; Sen et al., 2005). EM4 cells are human embryonic kidney 293 cells modified to increase their adherence to tissue culture plastic (Robbins and Horlick, 1998).

### Mice

All experiments were reviewed and approved by the Columbia University Medical Center’s Institutional Animal Care and Use Committee (IACUC). Mice were bred and housed in facilities at the William Black Medical Research Building. Same-sex animals of mixed genotypes are housed four to five per cage in a humidity- and temperature-controlled room, and mice are provided food and water *ad libitum*. Mice are maintained on a 12 h light/dark cycle (lights on at 7:00 A.M.).

### Mouse lines

#### Conditional Flotillin-1 alleles (Flot1 flox/flox)

*Flot1^flox/+^* (strain F1 CBAxC57Bl6/J) mice were generated via homologous recombination at the University of Connecticut Health Center’s Gene Targeting and Transgenic Facility. The targeting vector was designed with LoxP sites flanking Flot1 at exons 3 – 8. C57Bl/6 and CBA F1 hybrid ES cell lines were used for targeting. Germline chimeras were crossed with ROSA-26Flpe mice to remove the PGKneo cassette used for *in vitro* selection. Germline-positive F1 mice were backcrossed with the chimeric mice. Pups from this backcross were genotyped in order to identify loss-of-wildtype allele and confirm proper targeting. The homozygous Flot1 flox/flox line was maintained by consistently breeding littermates every 5-6 months.

#### DAT^iresCre^ (DIC)

DAT^iresCre^ (DIC, strain Bl6/J) mice were acquired from the Jackson Laboratory (Bar Harbor, Maine) (Backman et al., 2006). This line, in which DAT mRNA has an internal ribosome entry site (IRES) Cre knock-in at the 3’ UTR, restricts Cre-mediated recombination events to DAergic neurons. DIC mice were maintained as heterozygotes on a C57Bl/6 background strain by breeding with wild type (WT) C57Bl/6 mice every 5-6 months.

#### Flotillin-1 conditional knockout (Flot1 cKO)

The Flot1 cKO mice used in this study were derived from mating *Flot1^flox/flox^* mice with mice heterozygous for the DIC allele to create the F1 generation containing Flot1^fl/+^::DIC. For strain purposes, these Flot1^fl/+^::DIC mice were then crossed back with Flot1^flox/flox^ mice to generate the following littermates used in the experiments: Flot1^fl/fl^, Flot1(DAT) cHet, and Flot1(DAT) cKO.

#### Hprt^Cre/+^

Hprt^Cre/+^ mice (strain: Bl6/J) were obtained from the Jackson Laboratory (JAX no. 004302, Bar Harbor, Maine) (Tang et al., 2002), and backcrossed 10+ generations to create a Bl6/J line. The Hprt locus is located on the X chromosome and Cre expression results in 100% deletion of a conditional allele in oocytes. Hprt^Cre/+^ mice were maintained as heterozygotes on a C57Bl/6 background strain by breeding with WT C57Bl/6 mice every 5-6 months.

#### Flotillin-1 constitutive knockout (Flot1 KO)

Flot1^flox/flox^ males were crossed with heterozygous Hprt^Cre/+^ females to create Flot1^Δ/+^ mice. Flot1^Δ/+^ mice that did not carry Hprt^Cre/+^ were crossed together to generate the following experimental littermates: WT, Het, and KO.

#### Tamoxifen-inducible Cre (Actin^CreERTM/+^)

Actin^CreERTM/+^ mice (strain: Bl6/J) were obtained from the Jackson Laboratory (JAX no. 004682, Bar Harbor, Maine) (Hayashi and McMahon, 2002). This line expresses a tamoxifen-inducible Cre recombinase protein driven by a chicken beta actin promoter. The Cre recombinase is fused to a synthetic estrogen receptor, in which the ligand binding domain is modified, and is restricted to the cytoplasm. Upon tamoxifen injection, the estrogen receptor translocates into the nucleus, where excision of LoxP-flanked sequences occur. Actin^CreERTM/+^ mice were maintained as heterozygotes on a C57Bl/6 background strain by breeding with WT C57Bl/6 mice every 3-4 months.

#### Flotillin-1 inducible knockout (Flot1 iKO)

Flot1^fl/fl^ mice were crossed with Actin^CreERTM/+^ mice to create mice heterozygous for the Flot1 floxed allele and containing the Actin^CreERTM/+^ allele (Flot1^fl/+^:: Actin^CreERTM/+^). These Flot1^fl/+^:: Actin^CreERTM/+^ mice were then crossed back with the original Flot1^fl/fl^ mice to generate the following experimental littermates: Flot1^fl/fl^, iHet, and iKO.

### PCR Genotyping

#### DNA isolation

21-day-old mice were ear punched for identification and genotyping. Mice who have been retired from experiments were also re-genotyped by tail-clipping. The ear punches or tails were incubated at 55°C overnight in 0.5 mg/mL proteinase K made into 500 μL lysis buffer (50 mM Tris-HCl pH 8.0, 100 mM EDTA pH 8.0, 100 mM NaCl, 1% SDS). DNA was extracted by adding 700 μL phenol/chloroform/isoamyl alcohol (Fisher Scientific) to the digested sample, which was then centrifuged at 14,000 rpm for 5 min. To precipitate the DNA, 400 μL of the aqueous phase was carefully removed, mixed with 1 mL 100% ethanol by inverting tube, and spun at 14,000 rpm for 5 min. Once the supernatant has been decanted, the pellet was washed with 800 μL 80% ethanol and spun at 14,000 rpm for 5 min. The supernatant was removed and the DNA pellet was air-dried for 5 min at room temperature. Samples are resuspended in 50 μL (ear punch) or 200 μL (tail) of 10 mM Tris-HCl pH 8.0.

#### PCR

Each PCR reaction is 20 μL, consisting of 1 μL genomic DNA, 10 μM primer, 8 μL 5 PRIME HotMasterMix (5 PRIME GmbH), and water up to volume. The HotMasterMix contains Taq DNA polymerase, MgCl_2_, and dNTPs. Genomic DNA was extracted and the Flot1 LoxP sites were identified using the following primers: Flot1 LoxP forward: 5’**–**CTTCCCAGCCCTTACGTTCT**–**3’ and Flot1 LoxP reverse: 5’**–**GGGGTGGGGAGAATTCTATG**–**3’ (annealing temperature: 53°C); Frt forward: 5’**–** AGCGGATCCCATTACAGATG**–**3’ and Frt reverse: 5’**–**TCCCATGAGGGGATTACAAG**–**3’ (annealing temperature 57.5°C). Genotyping for the *DAT^iresCre^* allele was performed using the following primers: WT forward: 5’**–**TGGCTGTTGGTGTAAAGTGG**–**3’, WT reverse: 5’**–**GGACAGGGACATG GTTGACT**–**3’, and *DAT^iresCre^*: 5’**–**CCAAAAGACGGCAATATGGT**–**3’ (annealing temperature 62°C). Genotyping for the Hprt^Cre/+^ allele was performed using the following primers: WT forward: 5’**–** CACAGTAGCTCTTCAGTCTGATAAAA**–**3’, WT reverse: 5’**–**TTTCTATAGGACTGAAAGACTTG CTC**–**3’, Hprt^Cre/+^ forward: 5’**–**GCGGTCTGGCAGTAAAAACTATC**–**3’, and Hprt^Cre/+^ reverse: 5’**–**GT GAAACAGCATTGCTGTCACTT**–**3’ (touchdown PCR). Genotyping for the Actin^CreERTM/+^ allele was performed using the following primers: internal control forward: 5’**–**CTAGGCCACAGAATTGAAAGATCT**–**3’, internal control reverse: 5’**–**GTAGGTGGAAATTCTAGCATCATCC**–**3’, Actin^CreERTM/+^ forward: 5’**–**GCGGTCTGGCAGTAAAAACTATC**–**3’, and Actin^CreERTM/+^ reverse: 5’**–** GTGAAACAGCATTGCTGTCACTT**–**3’ (annealing temperature 51.7°C). To genotype for *Flot1* excision, Flot1 LoxP forward and Frt reverse were used at an annealing temperature of 57°C. All genotyping were done with a standard PCR protocol, unless otherwise stated.

### Drug Preparation and Administration

#### Amphetamine

Amphetamine (Sigma) was prepared in 0.9% saline at either 2.5 mg/kg or 5 mg/kg body weight (BW) for the open field behavior test. 5 mg/kg AMPH was used for animals prepared for biochemical assays.

#### Cocaine

Cocaine (Sigma) was prepared in 0.9% saline at 10 mg/kg BW for the open field behavior test and at 20 mg/kg BW for sucrose density gradients on mouse brains.

#### Tamoxifen

For temporal deletion of Flot1, experimental littermates for the Flot1 iKO were injected with 1.5 mg/ 10 g BW tamoxifen (Sigma) at 4 weeks of age. The tamoxifen solution was prepared into corn oil and 10% ethanol, which was heated at 37°C for 4 hours with vortexing every 30 min. Tamoxifen was administered to the iKO mice for five consecutive days. These mice were tested in open field behavior one month following their last injection when they are nine weeks old. All drugs were administered intraperitoneally (i.p.).

### Mouse Behavioral Testing

#### Open Field

All mice intended for behavioral testing were relocated and housed in our Satellite Animal Facility, where the open field equipment is located, two weeks prior to testing so that they could acclimatize to a new environment. Mice continued to be maintained on a 12 h light/dark cycle, for which 7:00 to 19:00 is the light cycle.

Testing began at 12-16 weeks of age for the Flot1 cKO, 8-12 weeks for the Flot1 KO mice, and 9-11 weeks for the Flot1 iKO mice. Open field was conducted between 9:00 A.M. and 7:00 P.M. during the light cycle. All mice were habituated to the open field prior to drug testing, which comprised of exposure to the open field chambers (43.2 cm x 43.2 cm x 30.5 cm) (Med Associates) for at least one hour per day for two to three consecutive days. All mice were assigned a chamber and were placed in the respective chambers for every subsequent run. For testing, mice were exposed to the open field for 1 hour, then given either saline, AMPH or COC (i.p.) then monitored for an additional 60 minutes. Each animal’s pathlength was collected and the data was binned into 5 min intervals.

### Electrochemistry

#### Fast scan cyclic voltammetry (FSCV)

FSCV was performed as previously described (Aguilar et al., 2017). Briefly, coronal brain slices with thickness of 250-300 µm containing cortex and striatum were prepared from 8-12 week old mice using a vibratome (Leica) and ice-cold cutting solution consisting of (in mM): glucose (10), NaCl (125.2), NaHCO_3_ (26), KCl (2.5), MgSO_4_ (3.7), NaH_2_PO_4_**·**6H_2_O (0.3), KH_2_PO_4_**·**6H_2_O (0.3). Slices were allowed to recover in the solution for 30 minutes at 34°C, and then transferred to recording ACSF (10 mM glucose, 125.2 mM NaCl, 26 mM NaHCO_3_, 2.5 mM KCl, 1.3 mM MgSO_4_, 2.4 mM CaCl_2_, 0.3 mM NaH_2_PO_4_**·**6H_2_O, 0.3 KH_2_PO_4_**·**6H_2_O). The temperature of the recording chamber was maintained at 32°C (± 2°C). Electrochemical recordings were performed with freshly cut disk carbon fiber electrodes 5 μm in diameter. The electrode was inserted ∼50 μm into the dorsolateral striatum. For cyclic voltammetry, a triangular voltage wave (−400 to +900 mV at 280 V/s vs. Ag/AgCl) was applied to the electrode every 100 ms. Currents were evoked by local electrical stimulation with a tungsten electrode (World Precision Instruments) and stimuli (100-400 µA, 1 ms duration) were delivered every 2 min by an ISO-flex stimulus isolator (AMPI) and Master-8 pulse generator (AMPI). Evoked currents were recorded with an Axopatch 200B amplifier with a low-pass Bessel filter setting at 10 kHz, digitized at 25 kHz (ITC-18 board; InstruTech). Triangular wave generation and data acquisition were controlled by a personal computer running a locally written (Dr. E. Mosharov, Columbia University, New York, NY) Igor Pro program (WaveMetrics). For amphetamine (AMPH) experiments, 10 µM AMPH was perfused into the bath for 20 min. The changes in evoked dopamine (DA) release were normalized by the average value of the peaks before AMPH perfusion. AMPH-induced DA efflux was detected 8-15 min after AMPH perfusion and the maximal peak of DA efflux was obtained in the presence of AMPH for the analysis. Carbon fiber electrodes were calibrated with 1 µM DA before and after recording, and voltammetric current unit was converted into molarity by calibration value.

#### High speed chronoamperometry (HSCA)

Experiments were performed as previously described (Dadalko et al., 2015). Briefly, striatal slices (300 µm) were prepared with a vibratome (Leica) in an ice cold oxygenated (95% O_2_/5% CO_2_) sucrose cutting solution consisting of 210 mM sucrose, 20 mM NaCl, 2.5 mM KCl, 1 mM MgCl2, 1.2 mM NaH_2_PO_4_, 10 mM glucose, 26 mM NaHCO_3_. Slices were then transferred to oxygenated ACSF at 28 °C for a minimum of 1 h. The ACSF consisted of 125 mM NaCl, 2.5 mM KCl, 1 mM MgCl_2_, 2 mM CaCl_2_, 1.2 mM NaH_2_PO_4_, 10 mM glucose, 26 mM NaHCO_3_, and 0.25 mM ascorbic acid. DA concentration was measured by chronoamperometry in slices as previously described (Gerhardt and Hoffman, 2001; Hoffman and Gerhardt, 1999). A carbon fiber electrode (100 μm length × 30 μm O.D.) coated with nafion for DA selectivity was positioned at a depth of 75-100 μm. The voltage was stepped from 0 mV to 550 mV for 100 ms and then back to 0 mV and the charging current of the microelectrodes was allowed to decay for 20 ms before the signals were integrated. Data were collected at a frequency of 1Hz with an Axopatch 200B amplifier. The integrated charge was converted to DA concentration based on in vitro calibration with DA. The oxidative signal measured in the slices is attributable to DA as the reduction/oxidation charge ratio is in the range 0.6-1.0.

### Immunohistochemistry

Staining was performed as previously described (Dragich et al., 2016; Yamamoto et al., 2000). Briefly, deeply anesthetized mice were transcardially perfused with 0.9% saline for 2 min followed by 4% paraformaldehyde (PFA) for 2 min. Brains were removed and post-fixed in 4% PFA for an hour before transfer to 30% sucrose in 1X phosphate buffer (P.B.) overnight. Post-fixed brains were fresh frozen in powdered dry ice, embedded with O.C.T. Compound (VWR), and stored at −80°C until use.

Immunohistochemistry (IHC) was performed on fixed sections. 30 μm free-floating coronal brain sections were cut into 1X P.B. containing 0.02% sodium azide. Sections were incubated in 10 mM sodium citrate, pH 9.0 for 30 min at 80°C for antigen retrieval. They were washed twice with 1X PBS containing 0.02% Triton X-100 (Tx-100), 10 min/wash. The endogenous peroxidases on the sections were blocked by incubation in 1% hydrogen peroxide (H_2_O_2_) diluted into 1X PBS containing 0.02% Tx-100 for 15 min, then washed three times with 1X PBS containing 0.02% Tx-100. After the washes, sections were blocked for 1 hr with 0.4% Tx-100 and incubated with antibodies overnight at room temperature.

On the following day, after three 10-min washes in 1X PBS, sections were incubated in biotinylated secondaries made into 0.4% Tx-100 for 1 hr. For sections stained with DAT, the secondary was anti-rat (1:2000) and for sections stained with TH, the secondary was anti-rabbit (1:2000). After three 10-min washes in 1X PBS, the signal was amplified using the standard VECTASTAIN Elite ABC HRP Kit (Vector Laboratories). The ABC complex was diluted into 1X PBS. Sections were placed in the diluted ABC solution for 1 hr at room temperature, followed by three 10-min washes in 1X PBS. Signal was detected using diaminobenzoate, or DAB (Sigma); 10 mg DAB was dissolved into every 20 mL 1X P.B. and activated with 0.00375% H_2_O_2_. Sections were washed three times in 1X P.B. then mounted on glass slides. The mounted sections were air-dried and dehydrated by dipping the slides in the order of the following solutions for 5 min each: 70% ethanol, 95% ethanol, 100% ethanol, and three times in xylene. The slides were coverslipped using Permount mounting media (Fisher Scientific).

### Quantification of DA levels

Quantification of DA and DOPAC levels was performed at the Neurochemistry Core at Vanderbilt University as described (Robertson et al., 2010). Dissected striata were homogenized in 100–750 µL of 0.1M TCA, containing 10-2 M sodium acetate, 10-4 M EDTA, 5ng/mL isoproterenol (as internal standard) and 10.5 % methanol (pH 3.8). Samples were spun in a microcentrifuge at 10,000 xg for 20 minutes. Supernatant were then analyzed for biogenic monoamines and/or amino acids using high pressure liquid chromatography with amperometric detection. Pellets were used to quantify protein. HPLC control and data acquisition are managed by Millennium 32 software.

### Sucrose density gradients

#### Striatal tissue preparation

Mice striata were prepared in 1% Surfact-Amps (Thermo Fisher), also known as 1% Brij58 with protease inhibitors (Pierce). Samples were homogenized and run through a 19-gauge needle 15 times followed by 25-gauge needle 15 times. The lysate was incubated on ice for 30 min, and spun at 850g for 10 min at 4°C. 1% Brij58 with protease inhibitors was added to 100 μg of the supernatant (sup) to make a total of 500 μL, which was mixed with an equivolume of 80% sucrose (wt/vol) in 150 mM NaCl. The mixture was transferred to an ultracentrifuge tube, which was then layered on with 1 ml of each of the following: 30% sucrose (wt/vol), 15% sucrose (wt/vol), and 5% sucrose (wt/vol). The tube was placed in a chilled SW60 Ti swinging-bucket rotor. Centrifugation was carried out in a Beckman ultracentrifuge at 43,200 rpm for 20 hrs at 4°C. Ten fractions of equal volumes were collected from top to bottom of the gradient. For the gradients in which mice were injected with drugs, mice were sacrificed 30 min after 5 mg/kg AMPH injection and 10 min after 20 mg/kg injection; both times are when mice have reached maximal hyperlocomotor behavior. The striata were processed as described.

To see whether cholesterol depletion would affect the partitioning of DAT in detergent-resistant membranes, 10 mM methyl-beta-cyclodextrin (MβC) was incubated with 100 μg striatal lysate for 40 min before being processed as described.

#### Striatal synaptosome preparation

cKO mice with Flot1^fl/fl^ and Flot1(DAT) cKO genotypes were sacrificed and their brains were removed into ice-cold 1X PBS and the striatum was dissected. The striatum was homogenized in Homogenization Buffer (H.B.) consisting of 0.32 M sucrose, 2 mM EDTA, 20 mM HEPES pH 7.3 with protease inhibitors using a glass dounce tissue grinder. Samples were spun at 1000 rpm for 10 min at 4°C. The sup was pipetted into a new tube, after which it was spun at 14,000 rpm for 20 minutes at 4°C. The sup was aspirated and the pellet resuspended in 1X PBS. The resuspended pellet was subjected to sucrose density gradients.

#### Cell culture

HEK293 cells were transfected with hDAT(WT) and hDAT(KA) were collected 48 hrs later in RIPA (50 mM Tris pH 7.5, 150 mM NaCl, 0.25% sodium deoxycholate, 1 mM EDTA, 1% NP-40, 1% Tx-100) with protease inhibitors. The lysate was incubated on ice for 30 min then spun at 1000 rpm for 10 min at 4°C. The sup was used for sucrose density gradients.

#### Western blots

The gradient fractions are run with 5 μg protein as lysate loading control. 42 μL fraction (maximum volume) was used. Western blot was run as described (Eenjes et al., 2016). Band intensities were quantified using ImageJ (NIH) and normalized to the lysate loading control.

### Limited proteolysis

All steps were performed on ice. Fractions 4 and 5 from the sucrose gradients described above were combined separately from fractions 9 and 10 so that the combined fractions contain the same concentration of the protein-of-interest. Protease solutions were always made fresh on the day of experiment; trypsin and papain were made into water and 2-Nitro-5-thiocyanatobenzoic acid (NTCB) was made into 100% ethanol. An aliquot was taken from the combined fraction and vehicle was added to the aliquot for the 0 min time point. Protease was added to the remaining combined fractions, mixed thoroughly, and quickly aliquoted into separate tubes. Protease inhibitor cocktail (for trypsin and papain) or β-mercaptoethanol (for NTCB) were added at the specified time point. SDS sample buffer was added and the samples were boiled then run on a western blot.

To ensure that the sucrose concentration of the fraction does not differentially affect protease cleavage capability, a gradient was run without any protein and 50 μg/mL BSA was spiked into the same gradient fractions mentioned above. Limited proteolysis was performed as described above and results were visualized by coomassie staining.

### Immunoprecipitation

HEK293 cells transfected with the hDAT constructs were lysed with RIPA and protease inhibitors. 100 μg protein was incubated with 1µL anti-PIP_2_ antibody (ENZO) for 30 min at 4°C with gentle shaking, then incubated in Protein G microbeads (Miltenyi) overnight at 4°C with shaking and processed the following day as described (Cremona et al., 2011).

#### Phosphorylation

EM4 cells were transfected with a mix of siFlot1. After 72 hours, cells were washed twice with PBS and once with pre-warmed Kreb’s buffer (130 mM NaCl, 1.3 mM KCl, 1.2 mM MgSO_4_, 1.2 mM KH_2_PO_4_, 2.2 mM CaCl_2_, 2 mM NaH_2_PO_4_, 10 mM HEPES pH 7.4, 10 mM glucose, 0.1 % BSA, 2 mM NaHCO_3_). They were treated with 1 μM PMA or vehicle (DMSO) in Kreb’s buffer with phosphatase inhibitors 1 mM sodium vanadate and 10 mM sodium fluoride for 30 min at 37°C and collected using 1% Brij58 with protease and phosphatase inhibitor cocktails. For pSer7 detection on western blot, an immunoprecipitation must first be carried out. Cell lysate was incubated overnight at 4°C with anti-GFP conjugated to magnetic beads (Miltenyi) and IP was performed as previously described (Cremona et al., 2011).

### Immunofluorescence

Cells were fixed with methanol for 10 min at room temperature. Cells were washed two times with PBS, incubated in 0.1% Tx-100 for 10 min, washed three times with PBS and blocked with 3% BSA in PBS. Cells were incubated in primary diluted in 0.01% Tx-100 overnight at 4°C. They were washed three times with PBS and incubated with Alexa Fluor secondary antibodies (Thermo Fisher) made into 1 μg/ml Hoechst solution (Life Technologies) for two hours at room temperature. Images were acquired using a Leica TCS SP2 confocal microscope at 63x magnification and the accompanying software package.

### Dissection and immunoblotting of substantia nigra

Mouse brains were placed in a pre-chilled brain matrix (Ted Pella), which allowed slices of 1 mm to be generated. A slice was taken from the midbrain, placed in cold PBS and the substantia nigra dissected. Tissue was prepared in 1% Brij58 with protease inhibitors. Protein concentration was determined using the DC Protein Assay (Bio-rad Laboratories) and equal amounts of protein were prepared for western blot as previously described (Eenjes et al., 2016).

### Statistical analyses and figure creation

Statistical analysis was performed using Statview 5.0 (SAS Institute, Cary, NC). Mean differences of normally distributed data were tested using analysis of variance (ANOVA) as described in the figure legends. When differences with greater than 90% were detected, *post hoc* analyses were performed using the Fisher’s protected least significant difference test. Non-normally distributed data was assessed using the nonparametric test, Mann-Whitney U. Complete F-statistics with p-values are presented in the figure legends, as well as n-values. Adobe Creative Suite 5 (Photoshop and Illustrator) were used to create all figures. ImageJ (NIH) was used for image quantification.

