## Supplemental Figures for "Membrane composition influences the conformation and function of the dopamine transporter *in vivo*"

Fong, W.M. et al. Figure S1

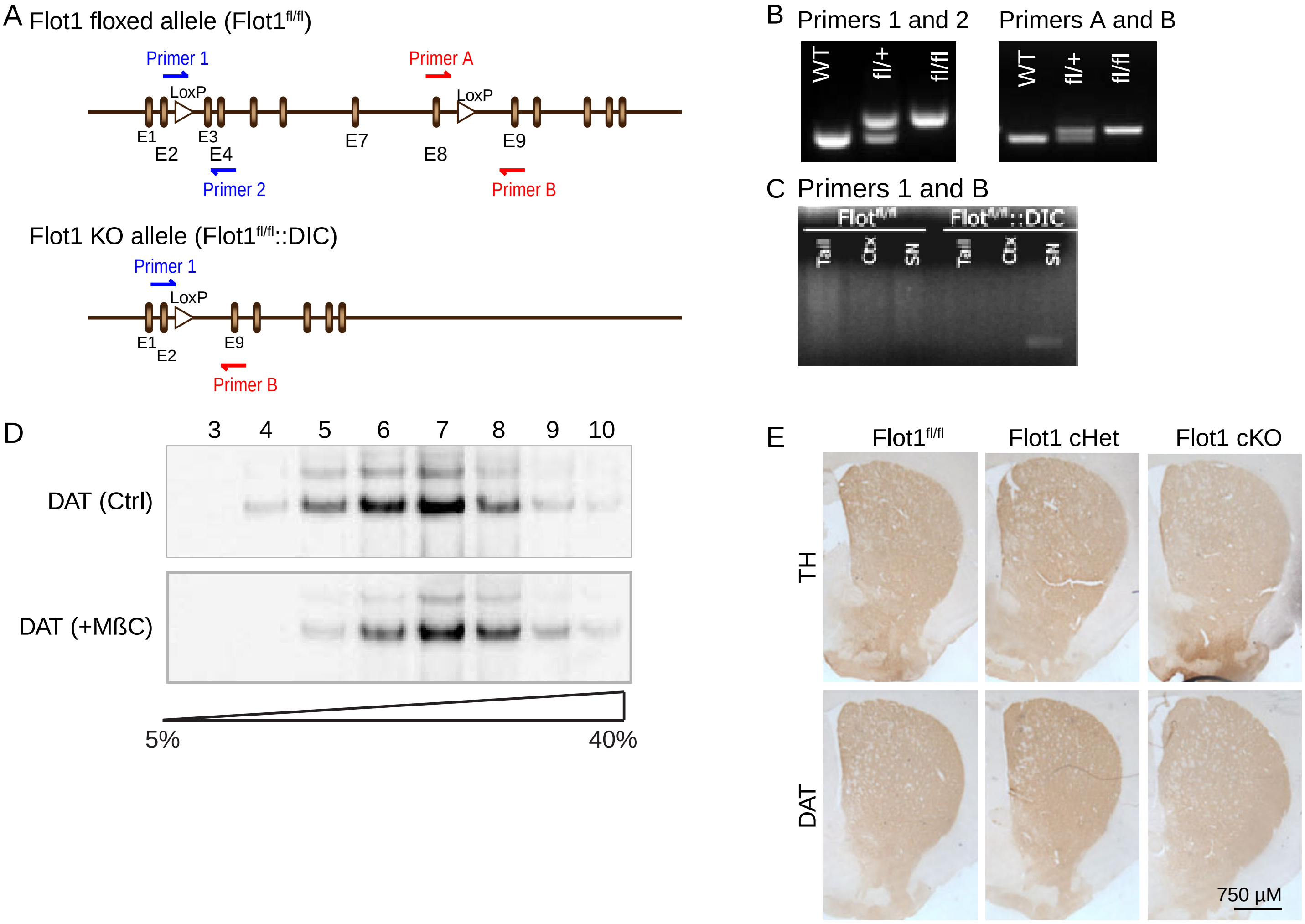


**Figure S1. The loss of Flot1 leads to no detectable changes in the DA system, but changes the partitioning of DAT into sterol- and sphingolipid enriched DRMs, as related to Figure 1.**

**A-C.** Design of the Flot1 conditional allele and resulting recombined allele of the mice in Fig. 1. **A**. LoxP sites were inserted between exons 2 and 3, and exons 8 and 9. PCR primers to detect the LoxP insertions are designated as Primers 1 and 2, and Primers A and B. **B**. PCR genotyping for the different primer pairs. **C**. PCR detection of recombined allele.The recombined Flot1 allele can be detected in DNA collected from substantia nigra (SN) of Flot1(DAT) cKO mice (Flot$^{fl/fl}$::DATiresCre (DIC)), but not in DNA from cortex (Ctx) or tail. Similarly, no recombination is detected in the Flot1$^{fl/fl}$ mice.

**D.** Sucrose density gradients of DAT striatal lysates (1% Brij58) from Flot1$^{fl/fl}$ mice in the absence (top) or presence (bottom) of methyl-beta-cyclodextrin (MßC). The chelation of cholesterol by MßC leads to the loss of DAT partitioning into the more buoyant fractions that are positive for Flot1 (See Figure 1A).

**E.** Immunohistochemistry against tyrosine hydroxylase (TH) and the dopamine transporter in Flot1$^{fl/fl}$, cHet and cKO striata. No detectable difference in staining was observed, similarly to staining in substantia nigra (See Figure 1B).

Fong, W.M. et al. Figure S2

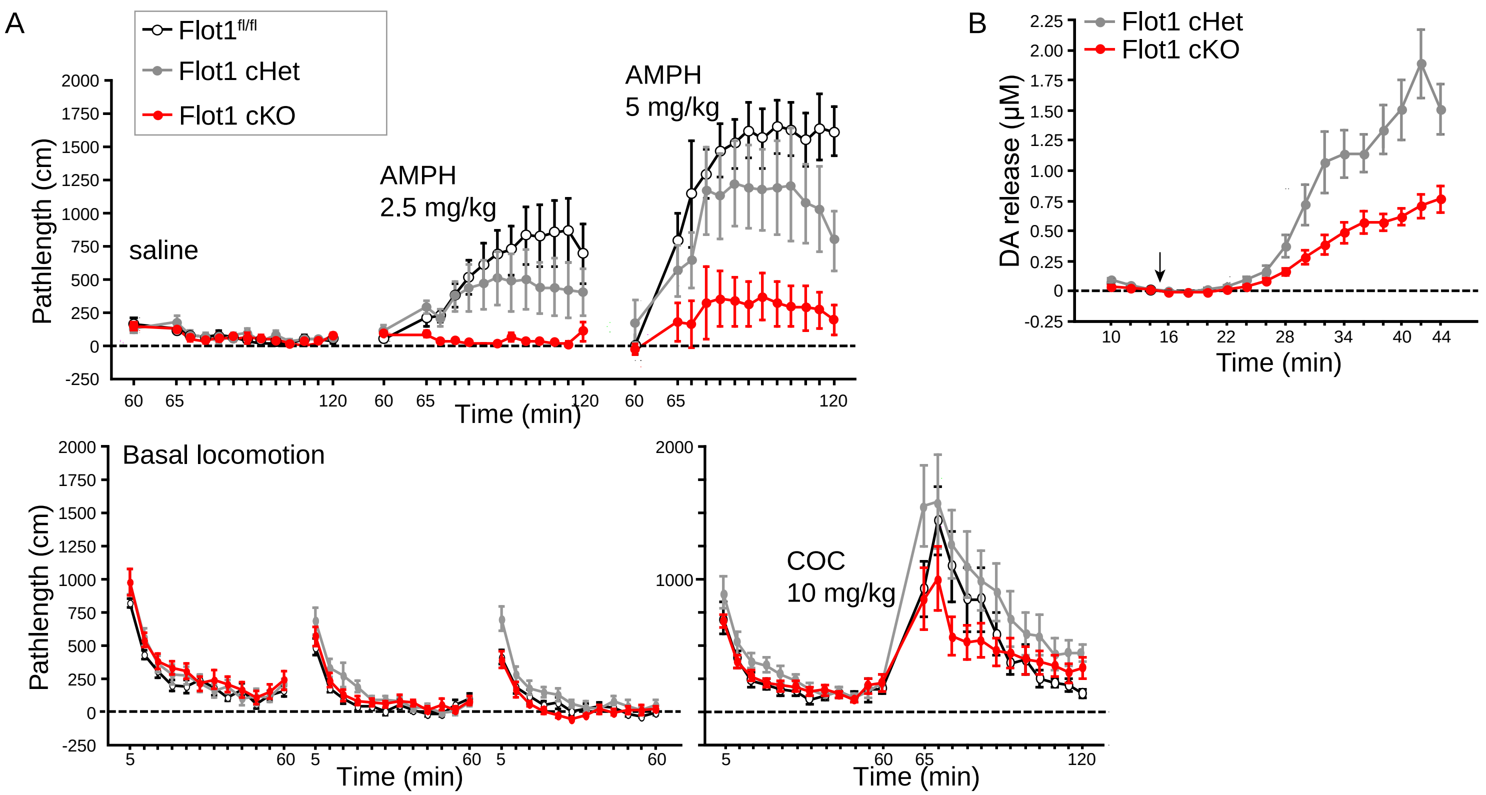


**Figure S2. Flot1(DAT) cKO mice show diminished AMPH-induced DA efflux through DAT, as related to Fig. 2.**

A**.** Locomotor response in female mice show a similar Flot1-dependence as males, with a diminished response to AMPH, and no difference in basal behavior or response to COC. Statistics (RM-ANOVA): ***Saline:*** Genotype effect ($F_{(2,33)}$=0.584, p=0.5635). ***AMPH 2.5 mg/kg***: Genotype effect ($F_{(2,33)}$=5.171, p=0.0111). Treatment (Rx) effect ($F_{(12,396)}$=6.958, p<0.0001). Interaction Genotype and Rx ($F_{(24,396)}$ =3.448, p<0.001). Fisher PLSD reveals a difference in Flot1 and cKO (p=0.0033), cHet and cKO (p=0.0472) but not Flot1 and cHet (p=0.2762). ***AMPH 5 mg/kg***: Genotype effect ($F_{(2,33)}$=8.031, p=0.0014). Rx effect ($F_{(12,396)}$=15.236, p<0.0001). Interaction Genotype and Rx ($F_{(24,396)}$= 1.969, p=0.0038). Fisher PLSD reveals a difference in Flot1 and cKO (p=0.0004), cHet and cKO (p=0.0144) but not Flot1 and cHet (p=0.1822). ***COC 10mg/kg:*** Genotype effect ($F_{(2,33)}$=2.136, p=0.1342). ***Basal:*** Day 1: Genotype effect ($F_{(2,33)}$=0.820, p=0.4493); Day 2: Genotype effect ($F_{(2,33)}$=1.828, p=0.1766); Day 3: Genotype effect ($F_{(2,33)}$=4.446, p=0.0195). Fisher PLSD reveals a difference in cHet and Flot1 (p=0.0370) and cKO (p=0.007), but not in Flot1 and cKO (p=0.4765). n=12 mice. Data shown as mean ±S.E.

**B*.*** Current amperometry in striatal slice preparations from the Flot1 cKO mice demontrate significantly diminished AMPH-induced release of DA through DAT. cHet are used as a control for the presence of $DAT^{iresCRE}$.Statistics (RM-ANOVA): Genotype effect ($F_{(1,16)}$=19.247, p=0.0005). n=3 cHet mice, 2 slices per mouse; n=3 cKO, 3 slices per mouse.

Fong, W.M. et al. Figure S3

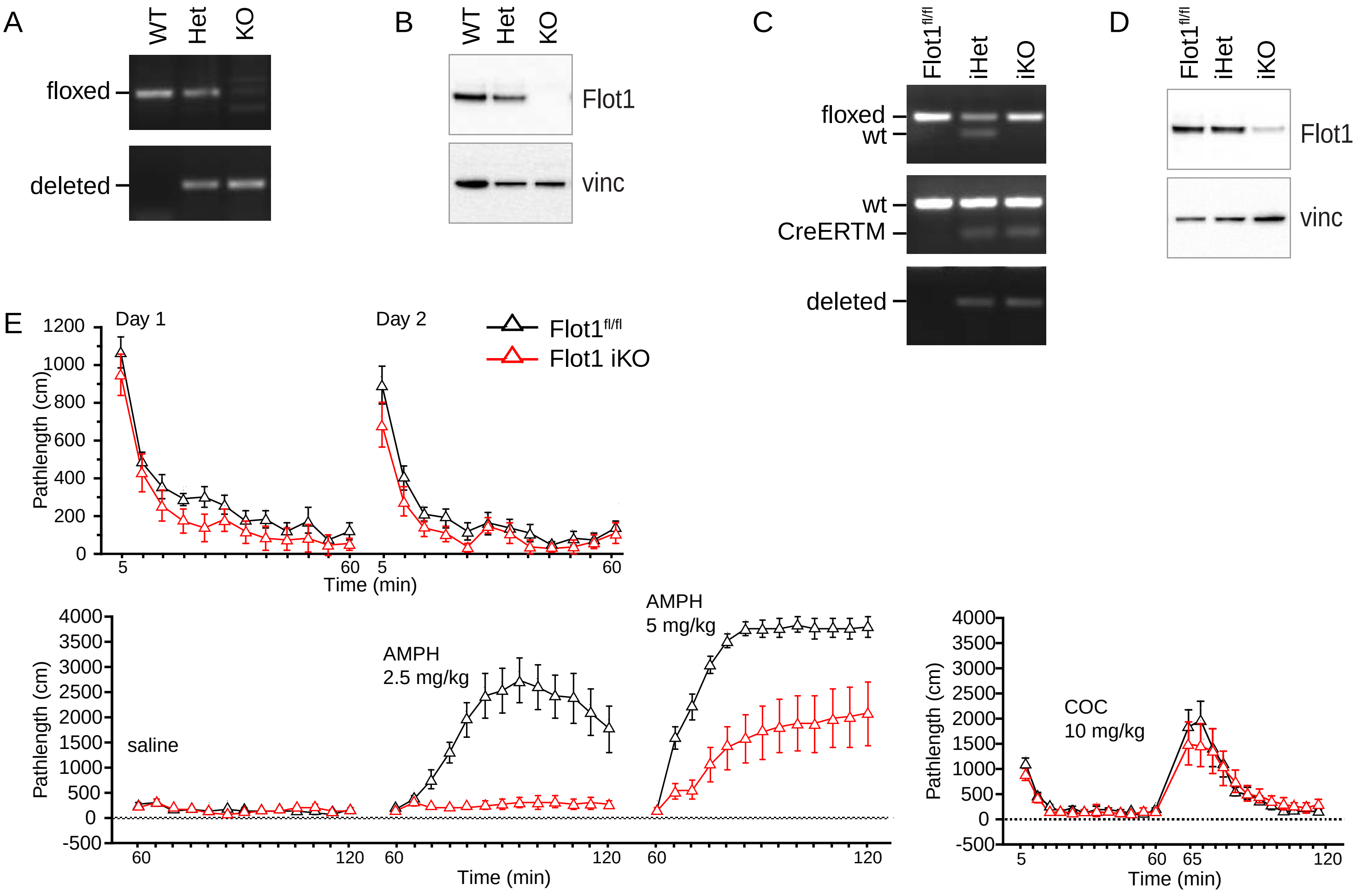


**Figure S3. Flot1 KO vs. Flot1 iKO mice reveal that compensatory events can mask the role of Flot1 in vivo, as related to Fig. 3.**

**A, B.** Flot1 KO mice. (**A**) PCR detection of Flot1 conditional allele versus deleted null allele in the WT, Het and KO mice. (**B**) Immunoblotting of whole brain lysates from WT, Het and KO mice.

**C, D.** Flot1 iKO mice. (**C**) PCR detection of Flot1 conditional allele versus WT allele (top), the presence of Actin CreERTM (middle) and the null allele (bottom) in $Flot1^{fl/fl}$, iHet and iKO mice. (**D**). Immunoblotting of tam-treated whole brain lysates from $Flot1^{fl/fl}$, iHet and iKO mice. n = 3.

**E.** Locomotor response in iKO female mice show similar Flot1-dependence as iKO males, with a diminished response to AMPH, and no difference in basal behavior or respose to COC. Statistics (RM-ANOVA): ***Basal:*** Day 1: Genotype effect ($F_{(2,25)}$=2.213, p=0.1304); Day 2: Genotype effect ($F_{(2,25)}$=1.598, p=0.2223). ***Saline:*** Genotype effect ($F_{(2,25)}$=0.680, p=0.5157). ***AMPH 2.5 mg/kg***: Genotype effect ($F_{(2,25)}$=19.139, p<0.0001). Treatment (Rx) effect ($F_{(12,300)}$=16.742, p<0.0001). Interaction Genotype and Rx ($F_{(24,300)}$ =10.635, p<0.0001). Fisher PLSD reveals a difference in Flot1 and iKO (p<0.0001), ***AMPH 5 mg/kg***: Genotype effect ($F_{(2,25)}$=5.716, p=0.0090). Rx effect ($F_{(12,300)}$=55.078, p<0.0001). Interaction Genotype and Rx ($F_{(24,300)}$= 2.178, p=0.0014). Fisher PLSD reveals a difference in Flot1 and iKO (p=0.0025). ***COC 10mg/kg:*** Genotype effect ($F_{(2,25)}$=0.112, p=0.8942). n=10,9 mice (Flot1, iKO). Data shown as mean ±S.E.

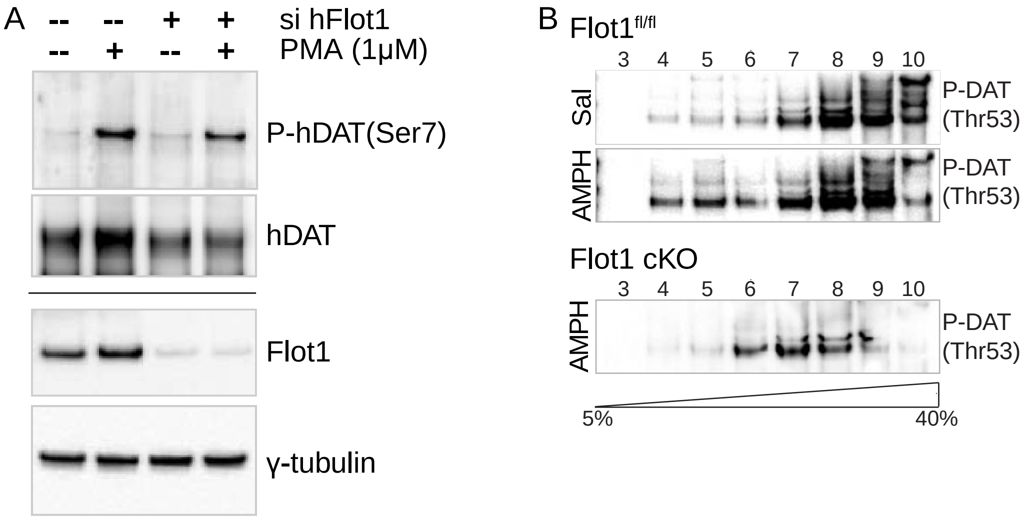


Figure S4. The partitioning of DAT into cholesterol rich membranes is not to  promote DAT phosphoryaltion.

A. hDAT(Ser7) is phosphorylated (P-hDAT(Ser7)) in response to PMA despite siRNA mediated depleation of Flot1 (si hFlot1). Experiments performed on stably expressed hDAT in EM4 cells. DAT is immunoprecipitated under the conditions shown. Western blotting by an antibody against P-hDAT(Ser7) reveals a basal level of phosphorylation, and that this phosphorylation is increased by the activation of PKC with PMA. Blots below show corresponding total lysates probed for Flot1 and Vinc as a loading control. n = 3.

B. Fractions from SDGs of 1% Brij 58 striatal lysates from Flot1$^{fl/fl}$ and Flot1 cKO mice, probed with an antibody raised against DAT(Thr53) (P-DAT(Thr53)) reveals that DAT(Thr53) is constitutively phosphorylated in both genotypes, and that the phosphorylated form is detected in all fractions.The corresponding immunoblot against DAT can be found in Fig. S4C top left for Flot1$^{fl/fl}$ saline, and Fig. S4A for Flot1$^{fl/fl}$ and Flot1 cKO AMPH. n = 2.

Fong, W.M. et al. Figure S5

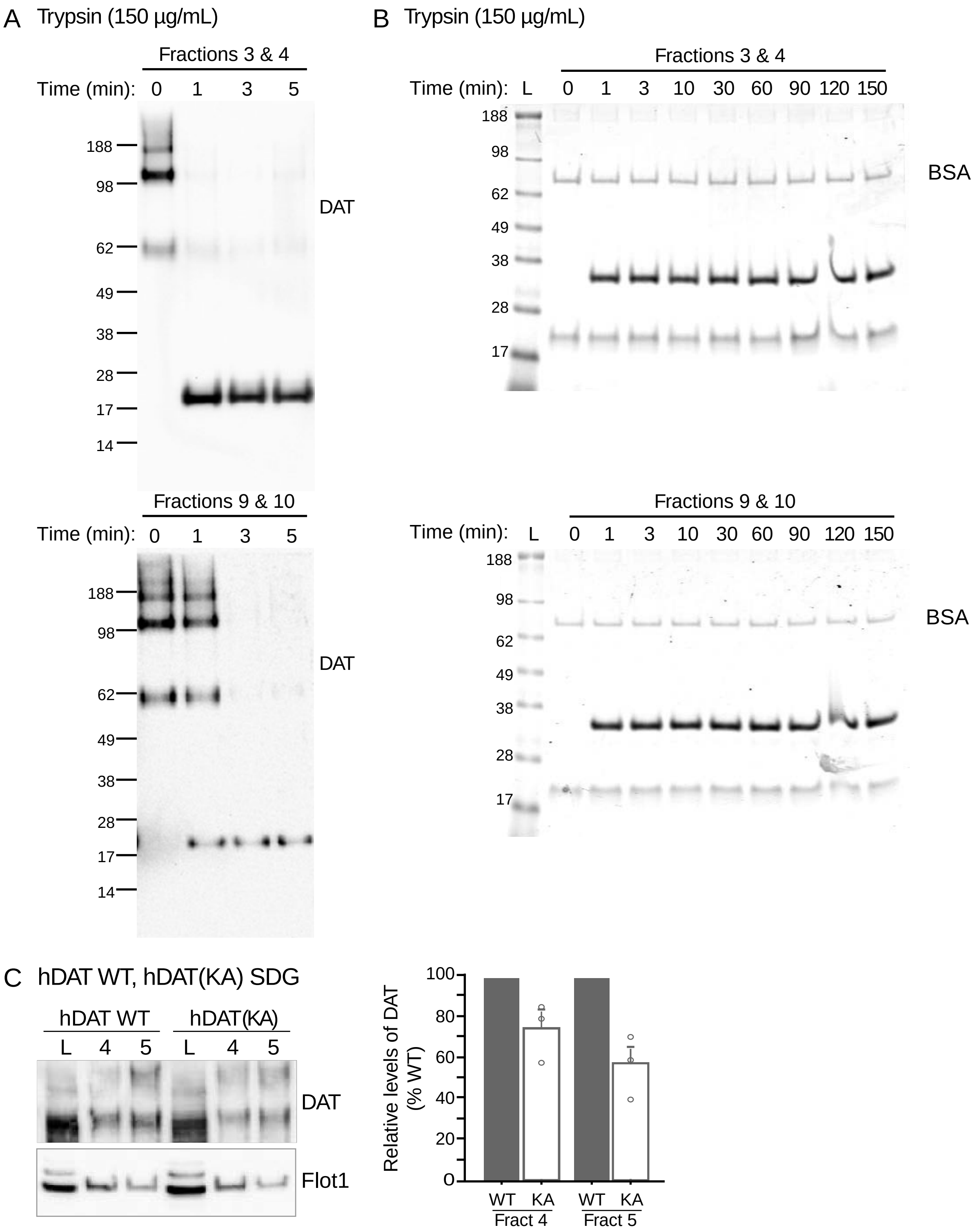


Figure S5. As related to Figure 5, proteolytic susceptibility of DAT differs between DRM and non-DRM fractions indicating DAT in DRMs are in a distinct conformation.

**A.** Proteolysis by 150 µg/mL of trypsin for the times indicated reveal that DAT in 1%Brij58 DRMs is more susceptible to cleavage than is DAT in non-DRM fractions.

**B.** BSA controls indicate that the different sucrose levels do not affect proteolytic rates as BSA is proteolyzed similarly in both lower and higher concentrations of sucrose in the buoyant and dense fractions respectively. The amount of proteolysis appears not to change over time, suggesting that trypsin works primarily in the first minute.

C. Diminished partitioning of hDAT(KA) upon SDG analyses of 1% Brij58 lysates. hDAT(KA) mutant continues to segregate to DRMs but to a lesser degree than hDAT WT (Mann-Whitney U test: Fraction 4, p=0.0495; Fraction 5, p=0.0495; n=3). This suggests that the diminishing the efficiency with which DAT interacts with PIP2 also affects its ability to remain within cholesterol-rich membranes, possibly through diminished interaction with Flot1.
